# A yeast optogenetic toolkit (yOTK) for gene expression control in *Saccharomyces cerevisiae*

**DOI:** 10.1101/663393

**Authors:** Jidapas (My) An-adirekkun, Cameron J. Stewart, Stephanie H. Geller, Michael T. Patel, Justin Melendez, Benjamin L. Oakes, Marcus B. Noyes, Megan N. McClean

## Abstract

Optogenetic tools for controlling gene expression are ideal for tuning synthetic biological networks due to the exquisite spatiotemporal control available with light. Here we develop an optogenetic system for gene expression control and integrate it with an existing yeast toolkit allowing for rapid, modular assembly of light-controlled circuits in the important chassis organism *Saccharomyces cerevisiae.* We reconstitute activity of a split synthetic zinc-finger transcription factor (TF) using light-induced dimerization. We optimize function of this split TF and demonstrate the utility of the toolkit workflow by assembling cassettes expressing the TF activation domain and DNA-binding domain at different levels. Utilizing this TF and a synthetic promoter we demonstrate that light-intensity and duty-cycle can be used to modulate gene expression over the range currently available from natural yeast promoters. This work allows for rapid generation and prototyping of optogenetic circuits to control gene expression in *Saccharomyces cerevisiae.*

---

The budding yeast *Saccharomyces cerevisiae* is an important chassis organism for synthetic biology and metabolic engineering [1, 2, 3]. These disciplines integrate biological parts (*e.g.* coding sequences, promoters) into biological circuits with novel cellular function. This process has become more routine in *Saccharomyces cerevisiae* due to the creation of large libraries of well-characterized parts [4, 5, 6, 7]. However, the inner workings of the cell continue to make the function and performance of engineered biological circuits unpredictable. Engineering efforts benefit greatly from the ability to tune the concentration of individual components to test and adjust circuit function. Additionally, tunability can allow circuits to be temporally and spatially adjusted to match dynamic constraints, such as the external environment or bioprocess phase [8, 9, 10].

Optogenetic approaches offer a potential solution for flexible tuning. Optogenetics take advantage of light-sensitive genetically encoded proteins to actuate processes inside of the cell in a light-dependent manner. Light is a powerful actuator as it inexpensive, easily controlled in time and space, and *Saccharomyces cerevisiae* contains no known native photoreceptors [11]. Here we report the construction and optimization of a light-activated transcription factor and associated components for use with an existing toolkit of yeast parts [4]. Addition of these optogenetic components to the toolkit allows for rapid and modular assembly of light-controlled circuits to tune gene expression dynamically and over the range defined by native yeast constitutive promoters.

We took advantage of the naturally occurring *Arabidopsis* cryptochrome CRY2 and blue-light dependent binding partner CIB1 to reconstitute the activity of a split transcription factor in a light-dependent manner. The feasibility of this approach was previously demonstrated by fusing CRY2 to the scGAL4 DNA-binding domain and CIB1 to the scGAL4 activation domain [12, 13, 14, 15]. The GAL4 protein is a native *S. cerevisiae* transcription factor and using the GAL4 DNA-binding domain (DBD) to create synthetic transcription factors leads to crosstalk with native GAL4-inducible promoters [16]. To avoid this crosstalk, we utilized a non-yeast DBD. The three-finger Zif268 DNA-binding domain from the Zif268 mouse transcription factor specifies a 9-bp sequence that occurs infrequently (<20) in the *S. cerevisiae* genome and this domain has been shown to be a powerful, orthogonal DNA-binding domain for generating chemically-inducible transcription factors in *Saccharomyces cerevisiae* [16, 17]. We fused the Zif268-DBD to the N-terminus of the CRY2 protein (ZDBD-CRY2) and the viral VP16 activation domain to CIB1 (VP16-CIB1) (**Figure 1A**). To create a promoter (pZF) responsive to our artificial transcription factor, we removed the GAL4 binding sites from the *sc*GAL1 promoter and integrated variable numbers and orientations of the Zif268 binding site (5’-GCG TGG GCG-3’) (**Supplemental Figure 1, 2**). Utilizing a promoter with four binding sites for the Zif268 DNA-binding domain (pZF(4BS)) upstream of yeVENUS and plasmids containing DBD-CRY2 and AD-CIB1 constructs we showed that the ZDBD-CRY2 based system induced gene expression as well as the original GAL4DBD-CRY2 system (Kennedy, *et al* 2010) (**Figure 1B**). Consistent with previous reports (Kennedy, *et al* 2010) a ZDBD-CRY2PHR construct containing only the CRY2 photolyase homology region (PHR) showed a stronger light-induced fold change as well as higher background gene expression **(Figure 1B)**. We verified that the GAL4-DBD did not exhibit crosstalk with the pZF promoters and that ZDBD-CRY2 did not activate expression from the scpGAL1 promoter (**Supplemental Figure 3**). We arrived at our final ZDBD-CRY2 construct and pZF promoters by testing different design considerations (*i.e.* linkers, nuclear localization signals, binding site number) as detailed in the **Supplemental Material** (**Supplemental Figures 2-5**). We domesticated our optogenetic components to interface with an existing yeast toolkit by removing the restriction enzyme sites (BsaI, BsmBI, NotI) used for the MoClo (modular cloning) assembly scheme [4, 18]. In the MoClo system, parts are categorized as “types” based on their function and location in the completed circuit (*e.g.* promoter types, coding sequence types) (**Figure 2A**). Our ZDBD-CRY2 and VP16-CIB1 components became coding sequence (“Type 3”) parts and the pZF promoters became promoter (“Type 2”) parts. All parts created in this study that are compatible with the Lee, *et al* [4] yeast toolkit are shown in **Supplemental Figure 6.** Using the modular cloning scheme with our optogenetic parts and additional parts from the yeast toolkit [4] allows for rapid assembly of multigene integration vectors containing all of the necessary components for light-induced expression of a gene of interest.

**Figure 1.**
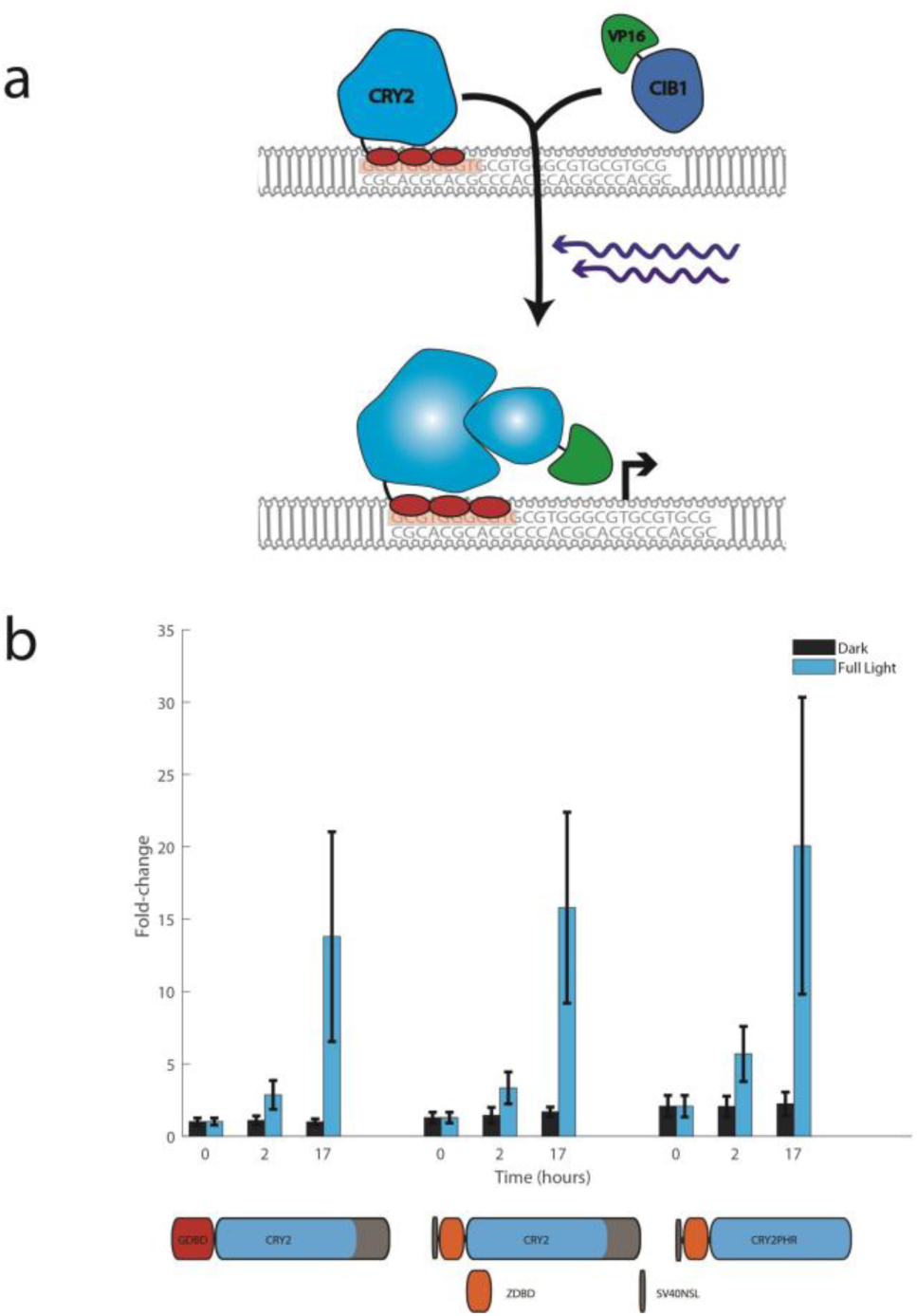
ZDBD-CRY2 AD-CIB1 optogenetic system: (A) Schematic of the ZDBD-CRY2 and CIB1-AD optogenetic system. In response to blue light, CRY2 undergoes a conformational change that allows CIB1 to bind CRY2. This recruits the activation domain to a promoter containing Zif268 binding sites (GCG-TGG-GCG). (B) Yeast cells transformed with the DBD-CRY2, and CIB1-AD plasmids and appropriate reporters (pGAL1-Venus or pZF(4BS)-Venus) were grown for up to 17 hours in 3mW/cm^2^ 460nm blue-light. Controls were grown in identical conditions without illumination. Induction is displayed as fold-change relative to the GAL4DBD-CRY2/CIB1-GAL4AD split transcription factor. Error bars indicate standard error of the mean.

**Figure 2.**
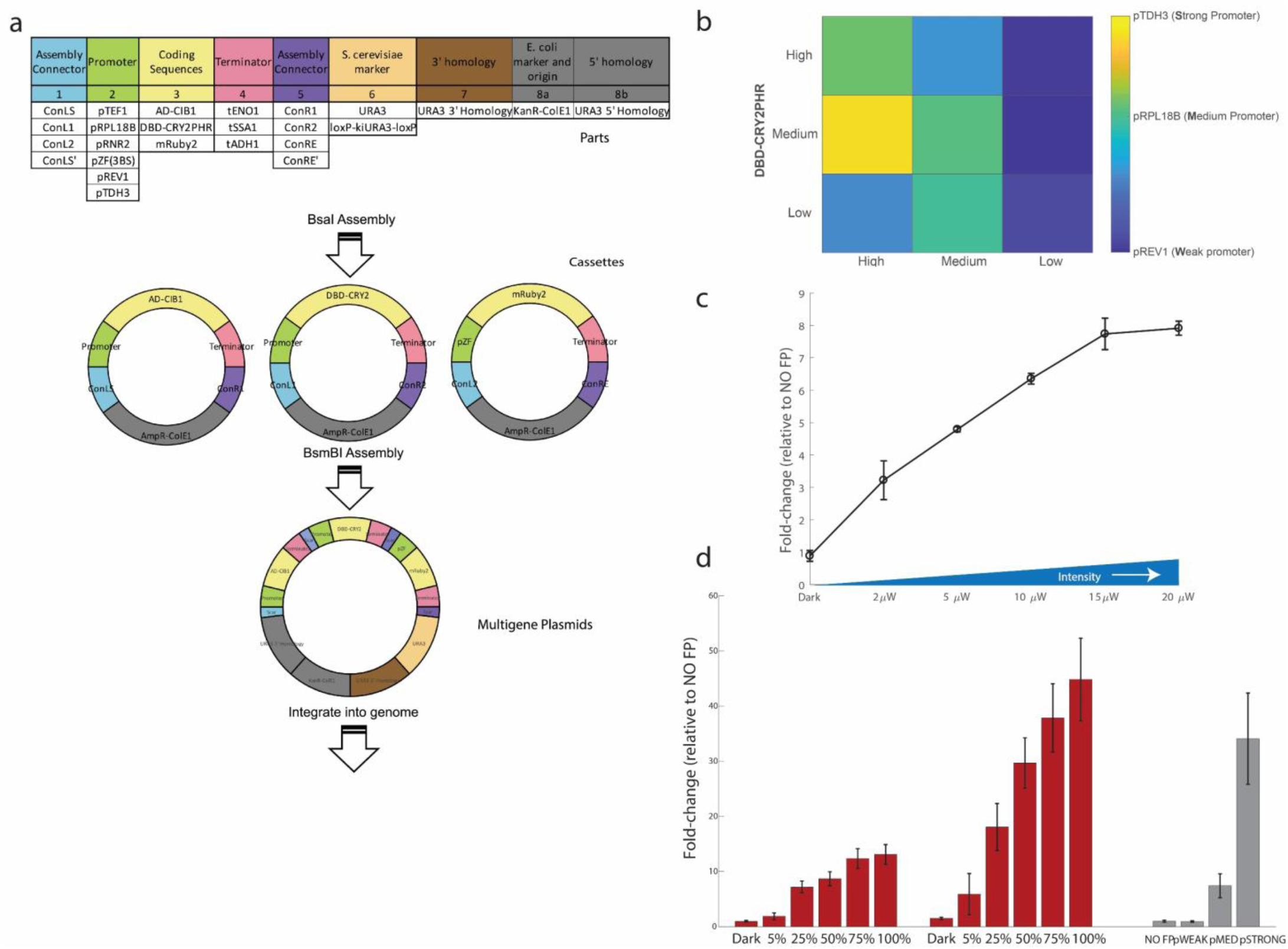
Optimization and Tuning of Light-Induced Gene Expression: (A) Schematic of circuit construction using the Yeast Toolkit scheme. Part plasmids contain unique upstream and downstream BsaI-generated overhangs to assemble into the appropriate position in “Cassette” plasmids. Cassette plasmids are fully functional transcriptional units that are further assembled into multigene plasmids using BsmBI assembly and appropriate Assembly Connectors. Figure adapted from Lee, *et al* 2015. (B) Fold-induction in gene expression in response to blue-light from pZF(3BS)-mRUBY2 was measured in nine strains expressing different ratios of the DNA-binding domain (ZDBD-CRY2PHR) and the activation domain (VP16-CIB1). Gene expression was compared to yeast strains expressing mRUBY2 under constitutive promoters of different strengths (pTDH3, pRPL18B, pREV1). (C) Gene expression in the ZDBD-CRY2_medium_/VP16-CIB1_medium_ optogenetic strain is tunable as a function of blue-light intensity. (D) Gene expression in the ZDBD-CRY2_medium_/VP16-CIB1_medium_ and the ZDBD-CRY2_medium_/VP16-CIB1_strong_ optogenetic strain is tunable as a function of light duty cycle. Fluorescence was measured as fold-change relative to the non-fluorescence control strain which contains a spacer region in place of pZF-mRUBY2 (yMM1477).

We demonstrated the utility of the toolkit workflow (**Figure 2A**) by utilizing it to optimize our synthetic split TF. We reasoned that the concentration and ratio of the two TF components, the DNA-binding component (ZDBD-CRY2PHR) and the activation component (VP16-CIB1), might be important for circuit function. We generated nine different optogenetic constructs with the two components under low, medium, and high-strength yeast promoters. To test gene expression control, these integrating vectors also contained mRUBY2 under the control of the pZF(3BS) promoter. We found that expression of AD-CIB1 under a high strength promoter (pTDH3) and DBD-CRY2PHR under a medium strength promoter (pRPL18B) gave us maximal expression from pZF(BS) (**Figure 2B, Supplemental Figure 7A**). However, this ZDBD-CRY2_medium_/VP16-CIB1_strong_ strain exhibited a growth defect (**Supplemental Figure 7B,C**). The ZDBD-CRY2_medium_/VP16-CIB1_medium_ strain on the other hand, with both components under the control of pRPL18B, exhibited expression from pZF(3BS) equivalent to a medium strength yeast promoter without exhibiting growth defects (**Figure 2B, Supplemental Figure 7**). The dosage of the DBD-CRY2PHR and AD-CIB1 components also affected the background (dark) induction (**Supplemental Figure 7A**) as well as the speed of induction after initial light exposure (**Supplemental Figure 8**). This was a crude dialing of protein concentration but suggests that one way to further optimize a split TF system would be by carefully titrating the total and relative dosage of each component, a knob not available in single-component and homogeneous two-component optogenetic systems [8, 19, 20].

One of the advantages of light as an inducer is the ability to tune its intensity and duty cycle in cultures of cells. We examined our ability to tune output from the ZDBD-CRY2_medium_/VP16-CIB1_medium_ strain as a function of light intensity. We found that we could tune output from the ZDBD-CRY2_medium_/VP16-CIB1_medium_ system up to 15µW/cm^2^ of light, at which point output from the system saturated (**Figure 2C)**. We also measured output from the ZDBD-CRY2_medium_/VP16-CIB1_medium_ and ZDBD-CRY2_medium_/VP16-CIB1_strong_ strains as a function of duty cycle. We varied light at 15µW/cm^2^ from a duty cycle of 5% (1min on/19 min off) to 100% (constant light). Gene expression output increased as a function of duty cycle, and by utilizing either the ZDBD-CRY2_medium_/VP16-CIB1_medium_ or ZDBD-CRY2_medium_/VP16-CIB1_strong_ strain we could achieve gene expression outputs equivalent to the weakest and strongest promoters in the Lee, *et al* yeast toolkit (**Figure 2D**). Thus, by putting the expression of a circuit component under the control of pZF and using light chemostats or programmable LED plates [21, 22, 9] one could continuously and dynamically tune component concentration and monitor its effect on circuit function. To allow this optogenetic machinery to be easily integrated into a yeast strain of interest we created a yeast marker (“Type 6”) part containing kiURA3 flanked by loxP sites to allow for marker recycling. We created a pre-assembled integration vector (**Supplemental Figure 6**) and used it to integrate ZDBD-CRY2_medium_/VP16-CIB1_medium_ and pZF(3BS)-mRUBY2. We verified that integration of the ZDBD-CRY2_medium_/VP16-CIB1_medium_ with a standard scURA3 marker or the kiURA3 marker before and after loxing by CRE-recombinase expression did not affect light-induced expression of mRUBY2.

Optogenetic approaches for controlling natural and synthetic biological networks are garnering attention as the toolkit expands and more powerful applications are demonstrated [22, 8]. Here we report an orthogonal light-inducible transcriptional activator for gene expression control in *S. cerevisiae.* We have engineered this transcription factor to be compatible with an existing yeast toolkit [4] which allows circuits for light-controlled gene expression to be assembled, integrated into the yeast genome, and tested in less than two weeks. We utilized this rapid prototyping to optimize the ratio and concentration of the two halves of our split transcription factor for maximal light-inducible gene expression with minimal growth defects. Both light intensity and duty cycle can be used to tune output from this gene expression system. We anticipate that this expansion of the yeast toolkit will be very useful to the community, as it will allow for rapid assembly of synthetic circuits (using the toolkit format) with one or more components that can be dynamically tuned with light. This will allow for circuit optimization as well as real-time light-based control of circuit output.

## Associated Content

Supporting Information is available online.

## Author Contributions

MNM, CS, MN, BO, MP, and MA designed experiments. BO, CS, MA, SG, and MP built constructs. SG, MA, MP, BO, and JM performed the experiments. MA, SG, MP, and MNM analyzed data. MNM drafted the manuscript. All authors contributed to the discussion and editing of the manuscript.

### Notes

The authors declare no competing financial interests.

## Acknowledgements

The authors would like to acknowledge discussion and helpful comments from the members of the McClean lab throughout the project. We thank K. Sweeney and T. Scott for assistance with Matlab and Python code for analyzing flow cytometry and growth curve data. This work was supported by the National Institutes of Health [1R35GM128873] and a Lewis-Sigler Fellowship from Princeton University (M.N.M). Flow cytometry was enabled by the University of Wisconsin Carbone Cancer Center Support Grant P30 CA014520 as well as the Princeton Flow Cytometry Resource Facility. Megan Nicole McClean, Ph.D., holds a Career Award at the Scientific Interface from the Burroughs Wellcome Fund.

## References

1. N. A. Da Silva and S. Srikrishnan, “Introduction and expression of genes for metabolic engineering applications in Saccharomyces cerevisiae,” FEMS Yeast Research, pp. 197–214, 2012.

2. K.-K. Hong and J. Nielsen, “Metabolic engineering of Saccharomyces cerevisiae: a key cell factory platform for future biorefineries,” Cellular and Molecular Life Sciences, vol. 69, pp. 2671–2690, 2012.

3. K. Buchholz and J. Collins, “The roots-a short history of industrial microbiology and biotechnology,” Applied Microbiology and Biotechnology, vol. 97, pp. 3747–3762, 2013.

4. M. Lee, W. Deloache and D. a. J. D. Cervantes, “A highly characterized yeast toolkit for modular, multipart assembly,” ACS Synthetic Biology, vol. 4, no. 9, pp. 975–86, 2014.

5. A. A. Reider, L. d’Espaux, M. Wehrs, D. Sachs, R. Li, G. Tong, M. Garber, O. Nnadi, W. Zhuang, N. Hillson, J. Keasling and A. Mukhopadhyay, “A Cas9-based toolkit to program gene expression in Saccharomyces cerevisiae,” Nucleic Acids Research, vol. 45, no. 1, pp. 496–508, 2017.

6. B. Chen, H. Lee, Y. Heng, N. Chua, W. Teo, W. Choi, S. Leong, J. Foo and M. Chang, “Synthetic biology toolkits and applications in Saccharomyces cerevisiae,” Biotechnology Advances, vol. 36, no. 7, pp. 1870–1881, 2018.

7. M. Jensen and J. Keasling, “Recent applications of synthetic biology tools for yeast metabolic engineering,” FEMS Yeast Research, vol. 15, no. 1, pp. 1–10, 2015.

8. E. Zhao, Y. Zhang, J. Mehl, H. Park, M. Lalwani, J. Toettcher and J. Avalos, “Optogenetic regulation of engineered cellular metabolism for microbial chemical production,” Nature, vol. 555, no. 7698, pp. 683–687, 2018.

9. A. Milias-Argeitis, M. Rullan, S. K. Aoki, P. Buchmann and M. Khammash, “Automated optogenetic feedback control for precise and robust regulation of gene expression and cell growth,” Nature Communications, vol. 7, p. 12546, 2016.

10. T. D. Scott, K. Sweeney and M. N. McClean, “Biological signal generators: integrating synthetic biology tools and in silico control,” Current Opinion in Systems Biology, vol. 14, pp. 48–65, 2019.

11. F. Salinas, V. Rojas, V. Delgado, E. Agosin and L. F. Larrondo, “Optogenetic switches for light-controlled gene expression in yeast,” Applied microbiology and biotechnology, vol. 101, pp. 2629–2640, 2017.

12. S. Fields and O.-k. Song, “A novel genetic system to detect protein-protein interactions,” Nature, vol. 40, no. 3, pp. 245–246, 1989.

13. G. P. Pathak, D. Strickland, J. D. Vrana and C. L. Tucker, “Benchmarking of optical dimerizer systems,” ACS Synthetic Biology, vol. 3, no. 11, pp. 832–838, 2014.

14. S. Shimizu-Sato, E. Huq, J. Tepperman and P. Quail, “A light-switchable gene promoter system,” Nature Biotechnology, vol. 20, pp. 1041–1044, 2002.

15. M. Kennedy, R. Hughes, L. Peteys, J. Schwartz, M. Ehlers and C. Tucker, “Rapid blue-light-mediated induction of protein interactions in living cells,” Nature Methos, vol. 7, no. 12, pp. 973–975, 2010.

16. R. McIsaac, B. Oakes, X. D. K. Wang, D. Botstein and M. Noyes, “Synthetic gene expression systems with rapid, tunable single-gene specificity in yeast,” Nucleic Acids Research, vol. 41, no. 4, p. e57, 2013.

17. R. S. McIsaac, P. A. Gibney, S. S. Chandran, K. R. Benjamin and D. Botstein, “Synthetic biology tools for programming gene expression without nutritional perturbations in Saccharomyces cerevisiae,” Nucleic Acids Research, vol. 42, no. 6, p. e48, 2014.

18. E. Weber, C. Engler, R. Grutzner, S. Werner and S. and Marillonnet, “A modular cloning system for standardized assembly of multigene constructs,” PLoS One, vol. 6, p. e16765, 2011.

19. D. Benzinger and M. Khammash, “Pulsatile inputs achieve tunable attenuation of gene expression variability and graded multi-gene regulation,” Nature communications, vol. 9, no. 1, p. 3521, 2018.

20. L. B. Motta-Mena, A. Reade, M. J. Mallory, S. Glantz, O. D. Weiner, K. W. Lynch and K. H. Gardner, “An optogenetic gene expression system with rapid activation and deactivation kinetics,” Nature Chemical Biology, vol. 10, no. 3, pp. 196–202, 2014.

21. K. Gerhardt, E. Olson, S. Castillo-Hair, L. Hartsough, B. Landry, F. Ekness, R. Yokoo, E. Gomez, P. Ramakrishnan, J. Suh, D. Savage and J. Tabor, “An open-hardware platform for optogenetics and photobiology,” Scientific Reports, vol. 6, no. 35363, 2016.

22. J. Melendez, M. Patel, B. L. Oakes, P. Xu, P. Morton and M. N. McClean, “Real-time optogenetic control of intracellular protein concentration in microbial cell cultures,” Integrative Biology, vol. 6, no. 3, pp. 366–372, 2014.

