## Supplemental Material for "A yeast optogenetic toolkit (yOTK) for gene expression control in *Saccharomyces cerevisiae*"

### Methods

#### Strains and Culture Media

##### Yeast strains and growth media

Yeast strains used in this study are shown in **Supplemental Table 1**. For light induction experiments followed by fluorescence assays, yeast were grown in either Synthetic Complete media (6.7 g/L Yeast Nitrogen Base without amino acids-DOT Scientific, 2% v/v glucose, 1% v/v KS amino acid supplement without appropriate amino acids) or Low Fluorescence Media (LFM) (1.7 g/L Yeast Nitrogen Base without ammonium sulfate, amino acids, folic acid, riboflavin-MP Biomedicals 4030-512, 5 g/L Ammonium sulfate-MP Biomedicals 808211, 1% v/v KS supplement, 2% v/v glucose-Fisher Scientific). Non-integrating plasmids were maintained by growing yeast in media lacking the appropriate amino acids required for plasmid selection.

##### Yeast Transformation

Yeast transformation was accomplished using standard lithium-acetate transformation [1]. For integrating plasmids, the integration was validated using either colony PCR or, when colony PCR proved difficult, by PCR of genomic DNA. Genomic DNA was extracted using the Bust n' Grab protocol [2]. Primers used for validating integrations are listed in **Supplemental Table 2**. All transformants were checked for the petite phenotype by growth on YEP-glycerol (1% w/v Bacto-yeast extract-BD Biosciences 212750, 2% w/v Bacto-peptone-BD Biosciences 211677, 3% [v/v] glycerol-Fisher Bioreagents BP229-1, 2% w/v Bacto-agar-BD Biosciences #214030) [3]. Only strains deemed respiration competent by growth on YEP-glycerol were used for subsequent analysis.

##### Bacterial strains and growth media

*Escherichia coli* strain DH5α was used for all transformation and plasmid maintenance in this study. *E. coli* were made chemically competent following either the Inoue method [4] or using the Zymo Research Mix & Go! Protocol (Zymo Research T3002). *E. coli* were grown on LB agar (10% w/v Bacto-Tryptone, 5% w/v Bacto Yeast Extract, 5% w/v NaCl, 15% w/v Bacto Agar) or LB liquid media (10% w/v Bacto-Tryptone, 5% w/v Bacto Yeast Extract, 5% w/v NaCl). Appropriate antibiotics were used to select for and maintain plasmids. Antibiotic concentrations used in this study were as follows: LB+CARB agar 100 µg/mL carbenicillin, LB+CARB liquid media 50 µg/mL carbenicillin, 25µg/ml chloramphenicol, 50µg/ml kanamycin.

#### Blue light induction

Blue-light induction was accomplished by either (1) illuminating cultures grown in glass culture tubes on a roller drum or (2) in a Light Plate Apparatus (LPA) [5]. Light intensity was measured and validated using a standard photodiode power sensor and power meter (Thorlabs #S120VC, Thorlabs #PM100D). The LPA was calibrated as described in Sweeney, *et al* [6] so that consistent light doses could be delivered between LPAs and between experiments.

For induction on a roller drum, biological replicates were selected from single yeast colonies and grown in Synthetic Complete media (lacking appropriate amino acids to maintain plasmid selection) at 30°C in the dark to mid-log. Each culture was then split and half of the culture was placed on the outside lane of a roller drum at room temperature in a glass culture tube. The other half of the culture was put in a test tube wrapped in foil on the inner lane of the roller drum. Three LEDs outputting 460nm blue light (Sparkfun COM-08718) were placed at the three, nine, and twelve o'clock positions of the roller drum and turned on at T=0 (~3mW/cm$^2$). At each timepoint, 500µl of culture was sampled into ice-cold PBS + 0.1% Tween and immediately placed at 4°C until being analyzed by flow cytometry.

For induction in the Light Plate Apparatus, yeast were grown overnight (12-16 hours) in 1ml of Low Fluorescence Media or Synthetic Complete in each well of the 24-well plate (Artic White, AWLS-303008) in the Light Plate Apparatus, in a light-proof coffin shaker at 30°C with constant shaking (250RPM). One glass microbead (Fisher Scientific 11-312A 3mm or 11-312B 4mm) was added to each well of the plate to increase aeration. The plate was covered with a Breathe-Easy® sealing membrane. After overnight growth, cultures were diluted to $OD_{600}$ = 0.2 and grown in a new 24-well plate on the LPA for the time indicated at the indicated light intensity. Timepoints were taken by transferring 50 µL of culture from each well to a 96 well plate containing 150 µL of PBS+0.1% Tween-20 in each well.

The LPA is programmable, such that the timing, intensity, and duration of illumination in each well can be controlled [5]. For the duty cycle experiment, the LPA was programmed to alternate between light on and light off with the following programs: turn on for 1 min, off for 19 min; on for 5 min, off for 15 min; on for 10 min, off for 10 min; on for 15 min, off for 5 min.

#### Flow Cytometry

Gene expression in response to blue light was assayed using fluorescent reporters and flow cytometry. Flow cytometry was performed on either a BD Biosciences LRSII Flow Cytometer (488nm laser and 505LP dichroic filter) or an Attune NxT Flow Cytometer (ThermoFisher Scientific) with 561nm excitation laser and 620/15nm filter cube. For assaying mRuby2 fluorescence on the Attune, the voltages of the flow cytometer were calibrated using rainbow beads so that the medians of FSS, SSC, and mRuby2 fluorescence of the rainbow beads were within 10% difference. The flow cytometry data was then analyzed using FlowJo software.

All samples were prepared for flow cytometry by diluting yeast cell culture (250-500µl) into 800µl of ice-cold PBS + 0.1% Tween-20. Samples were kept on ice or at 4°C until being analyzed. Samples run on the LPA were measured without sonication. Samples grown in glass culture tubes were sonicated with 10 bursts of 0.5 seconds each once diluted in PBS and prior to flow cytometry.

#### Construction and Optimization of the ZDBD-CRY2/AD-CIB1 Optogenetic Split Transcription Factor

To compare DBD-CRY2/AD-CIB1 combinations, yeast strain yMM1146 (Matα trp1Δ63 leu2Δ1 ura3-52) was cotransformed with appropriate plasmid combinations (DBD-CRY2/AD-CIB1/Reporter plasmid) as outlined in detail below using standard lithium acetate transformation [1]. Transformants were selected on and cultured in SC media with appropriate amino acids left out (SC-ura-leu-trp) for plasmid maintenance. Various plasmids were created and tested to determine an optimal ZDBD-CRY2/AD-CIB1 system (**Supplemental Table 3,** see below). Function of these plasmid combinations was tested by assaying for blue-light induced gene expression in glass culture tubes as outlined above.

##### Plasmid Construction

*AD-CIB1*

The VP16-CIB1 plasmid (pMM281) was created using yeast homologous recombination [7]. The VP16 activation domain from the GEV artificial transcription factor [8] was amplified from yMM1008 genomic DNA using oMM400/401. The pMM159 plasmid was cut with KpnI, which digests the plasmid between the existing SV40NLS and the GAL4AD. The PCR product containing VP16 and the digested plasmid were co-transformed into yMM83 (Mat a his3Δ1 leu2Δ0 LYS2 met15Δ0 ura3Δ0) and positive transformants that repaired the lesion in the plasmid with the VP16 activation domain (replacing the GAL4AD) were selected for growth on SC-Leu media. Plasmids were prepped from yeast and verified by sequencing.

*ZDBD-CRY2 with different linkers between Zif268DBD and CRY2*

The original ZDBD-CRY2 plasmids (pMM282, pMM283, pMM284) were made by molecular cloning and assembly into the pRS414 vector (pMM006 CEN TRP1) [Error! Reference source not found.]. The ADH1 promoter from pMM160 was amplified using oMM456/457 and cloned between the NgoMIV and NotI sites in pRS414. The FLAG(3X)-NLS-Zif268 cassette was amplified from plasmid pMN8-FLAG(3x)-NLS-Zif-Nuclease (a gift from the Noyes lab) using oMM458 and oMM407, 408 or 409 (to achieve different linker lengths between Zif268 and CRY2) and cloned between the NotI and NheI sites. The CRY2-tADH1 region from pMM160 was amplified using oMM383/oMM384 and cloned between the NheI and SacI restriction sites.

*pZF promoters*

The binding site reporter plasmids (pMM285-290) were made using the engineered GAL1 promoter from McIsaac, *et al* 2012 [9]. This GAL1 promoter was engineered to have XbaI and NotI sites on either side of the three native GAL4 (5'-CGG-$N_{11}$-CCG-3') binding sites. This promoter was cloned into pMM8 (pRS416 [10]) using NheI and XmaI sites. The fluorescent protein yEVenus from pMM223 (pKT90 [11]) was cloned between the XmaI and AscI sites using oMM421/423. To replace the GAL4 binding sites with binding sites for the Zif268 zinc-finger DNA-binding domain, pMM301 was digested with XbaI and NotI, and the oligo pairs oMM413-420, oMM481-486 were annealed and ligated into the pMM301 backbone. Plasmid pMM301 is the original GAL1 promoter from McIsaac, *et al* 2012 [9] in front of yEVENUS.

*ZDBD-CRY2 with different Nuclear Localization Signals (NLS)*

Plasmids pMM313 and pMM314 utilizing the SV40NLS and the H2BNLS respectively were made by digesting out the FLAG(3X)-SV40NLS tag in pMM284 between KpnI and NotI and annealing and ligating in either the oMM566/567 or the oMM568/569 oligonucleotide pairs to reinsert a lone SV40NLS-linker or H2B NLS-linker.

*Control Plasmids*

Plasmids pMM315 (GAL4AD scLEU2 CEN) and pMM316 (GAL4BD scTRP1 CEN) were constructed by yeast homologous recombination to remove the CIB1 and CRY2 open reading frames, while preserving GAL4AD and GAL4BD, from pMM159 and pMM160, respectively. Plasmid pMM159 was digested with ClaI and transformed with the annealed oligonucleotide pair oMM547/548. Positive transformants were selected on SC-LEU, plasmids were recovered from these yeast and verified by sequencing, resulting in pMM315. Plasmid pMM160 was digested with SalI and transformed with the annealed oligonucleotide pair oMM549/550. Positive transformants were selected on SC-Trp, plasmids were recovered from these yeast and verified by sequencing, resulting in pMM316.

*ZDBD-CRY2 without N-terminal FLAG (3x)*

Plasmids pMM317 (SV40NLS-ZCRY2 (L3) scTRP1) and pMM320 (SV40NLS-ZCRY2PHR scTRP1) were created by using yeast homologous recombination to remove the FLAG(3X) tag in pMM284 and using yeast homologous recombination to truncate CRY2. Plasmid pMM317 was constructed by cutting pMM284 with NotI and co-transforming this cut plasmid with annealed and extended oMM551/552 into yMM83[Error! Reference source not found.]. Positive transformants were selected on SC-Trp, purified from yeast, and verified by sequencing. Plasmid pMM320 was constructed by digesting pMM284 with both SalI and NotI, and transforming this digested plasmid with annealed and extended oMM551/552 as well as oMM562/563

#### Domestication for the YTK

The parts added to the Yeast Toolkit [12] are shown in **Supplemental Figure 6.** Domestication of parts for the Yeast Toolkit [12] requires removal of BsaI, BsmBI, and NotI restriction sites. This was accomplished using the Q5 site-directed mutagenesis kit (New England Biolabs E0554S) and appropriate primers as indicated in **Supplemental Table 2** to introduce a synonymous mutation to remove the undesirable restriction enzyme site**.** Domestication was verified by Sanger sequencing. Domesticated parts were inserted into the part-plasmid backbone (yTK001) as described in Lee, *et al* using primers in **Supplemental Table 2.**

#### Golden Gate Assembly of Cassette and Multigene Plasmids

Cassette plasmids (consisting of transcriptional units, *i.e.* promoter-coding sequence-terminator) and multigene plasmids (consisting of multiple transcriptional units linked together through assembly connectors with appropriate homology to integrate into the yeast genome) were assembled using BsaI assembly or BsmBI assembly as outlined in Lee, *et al* [12].

NEB Golden Gate assembly mix (E1600) was used for BsaI assembly. The 10 µL Golden Gate reaction mixture consisted of 1 µL of NEB Golden Gate Buffer (10x), 0.5 µL NEB Golden Gate assembly mix, 20 fmol of each plasmid, and water. We found that using commercially available NEB Golden Gate assembly mixture, as opposed to using BsaI, T7 Ligase, and T4 Ligase buffer, increases the reaction efficiency greatly. For BsmBI assembly, the protocol was adapted from Lee, *et al* [12] and each 10 µL BsmBI reaction mixture consisted of 0.5 µL BsmBI, 0.5 µL T7 Ligase, 1 µL T4 Ligase buffer, 20 fmol of each plasmid, and water.

The thermocycler program was adapted from Lee et al. (2015) [12] and consisted of 20-30 cycles of digestion and ligation (2 min at 37-42°C; 5 min at 16°C) followed by a final digestion (55-60 °C) and a heat inactivation step (80°C for 10-20 min). For final cassettes with internal BsaI cut sites (i.e. integration vectors), the reaction was ended with ligation, and final digestion and inactivation steps were omitted.

5 µL of reaction mixture was then transformed into DH5α competent *E. coli* and plated on LB plates with appropriate antibiotics. Plasmids were then extracted, digested with BsmBI or NotI-HF as a first-pass test, and sequenced with appropriate primers for final verification. For both BsaI and BsmBI assemblies, the efficiencies were found to be at least 50%. However, final cassettes with internal BsaI cut sites have notably lower assembly efficiency.

#### Construction of the ZDBD-CRY2/AD-CIB1 Dosage Strains

To understand how the dosage and ratio of the DNA-binding domain (ZDBD-CRY2) and activation domain (AD-CIB1) components of the optogenetic system affected function, we constructed nine strains with different ratios of ZDBD-CRY2 and AD-CIB1. These strains are yMM1458-1466 and have the genotypes shown in **Supplemental Table 1.** To construct these strains, we made multigene cassettes containing pPROMOTER-VP16-CIB1-tTerminator-pPROMOTER-ZDBD-CRY2PHR-tTerminator-pZF-mRUBY2 (pMM637-645) designed to integrate at the scURA3 locus. These multigene cassettes were made using BsmBI Golden Gate assembly (as described above) and cassette plasmids containing the ZDBD-CRY2, AD-CIB1, and pZF-mRUBY2 elements with appropriate assembly connectors (pMM620-624, 628-629). As benchmarks for mRUBY2 expression we also made multigene plasmids containing pCONSTITUTIVE-mRUBY2 instead of the pZF-mRUBY2 (pMM625-pM627) and integrated them into yMM1146 to create yMM1472-1480. As a no fluorescence control we created a multigene plasmid with a spacer in place of pZF-mRUBY2 (pMM619) and integrated this into yMM1146 to create the no-fluorescence controls yMM1473 and yMM1477.

#### Growth Assays

Yeast colonies were inoculated in 1 mL of LFM or SC-URA in a 24-well plate (Corning #3370) and grown for 24hrs in a light-proof coffin shaker at 30°C with shaking (250rpm). One glass bead was added to each well and the plate was covered with Breathe-Easy® sealing membrane (Sigma-Aldrich) to minimize evaporation. On the next day, the culture was diluted in 1 mL of LFM to $OD_{600}$ = 0.2 in a 24 well plate with cover (with no glass beads added). The $OD_{600}$ of the culture was measured every 15 min for 18 hours using the TECAN plate reader (Tecan Infinite M1000). Cells were grown for 18 hours with continuous double orbital shaking (120 rpm) and $OD_{600}$ readings taken every 15 minutes. Four readings were taken for each well for every time point. Growth rate µ was determined for the culture in each well by fitting the log-transformed $OD_{600}$ readings to the modified Gompertz equation described in [13] using the Trust Region Reflective algorithm implemented in SciPy [14, 15]. We observed that in some cases the early time points were too dilute to give consistent readings in the Tecan so that rather than normalize by the starting reading as described in Zwietering, *et al* [13] we added a fourth parameter $N_0$ to represent the starting concentration and fit the equation y= $A$ exp(-exp($\mu$ * e * ($\lambda$ - t) / $A$ + 1) ) + log $N_0$ where y is the log-transformed $OD_{600}$ readings.

#### Recycling of loxP-flanked markers

In order to allow for marker recycling utilizing the Cre-loxP system we created a loxP-KlURA3-loxP part plasmid (pMM519). The loxP-KlURA3-loxP cassette was amplified from pMM326 (pUG72; Gueldener, *et al* 2002) using oMM991/992 and assembled into the part entry vector (pYTK001/pMM452) using a BsmBI Golden Gate reaction. This part was further assembled into a multigene cassette designed for integration at the scURA3 locus (pMM617). This cassette plasmid contains 3'- and 5'- homology to the scURA3 locus flanking loxP-KlURA3-loxP. The Type234 GFP dropout is flanked by the special assembly connectors conLS' and conRE'. This pre-assembled integration vector allows additional cassettes flanked by assembly connectors to be assembled into this vector using a BsmBI Golden Gate reaction.

We used a BsmBI Golden Gate reaction with cassette plasmids pMM617, pMM621 (pRPL18B-CIB1VP16), pMM623 (pRPL18B-ZCRY2PHR), and pMM624 (pZF(3BS)-mRuby2) to generate pMM647 (pRPL18B-CIB1VP16-tENO1 pRPL18B-ZCRY2PHR-tSSA1 pZF(3BS)-mRuby2-tADH1 loxP-KlURA3-loxP scURA3 3' homology-KanR-ColE1-scURA3 5' homology). This multigene cassette plasmid was linearized with NotI and transformed into yMM1146 using a standard lithium acetate protocol [1]. Positive transformants were selected on SC-URA and verified by colony PCR and sequencing. This generated yeast strain yMM1468. Yeast strain yMM1468 is identical to the ZDBD-$CRY2_{medium}$/VP16-$CIB1_{medium}$ strain yMM1462 except that yMM1468 has the KlURA3 marker flanked by loxP sites instead of the standard scURA3 marker.

Cre-mediated recombination to recycle the KlURA3 marker was accomplished by adapting the CRE recombinase-mediated excision protocol from Carter and Delneri [16]. yMM1468 was transformed with 0.25-0.5 µg of pMM296 (pSH65, pGAL1-CRE $Bleo^R$). These transformants were plated onto YPD and then replica plated onto selective media (YPD +10µg/ml phleomycin (InvivoGen)) after overnight growth. To express CRE and induce recombination phleomycin resistant colonies were selected and grown overnight in 3ml of YP-Raffinose (1% w/v yeast extract (BD Biosciences), 2% w/v Bacto-peptone (BD Biosciences), and 2% w/v raffinose (Becton Dickinson 217410)). The following day, cells were harvested by centrifuging at 3750 rpm for 5 minutes, washed in sterile miliQ water, and resuspended in 10ml of YP-Galactose (1% w/v yeast extract (BD Biosciences), 2% w/v Bacto-peptone (BD Biosciences), 2% w/v galactose (BD Biosciences 216310)) at an $OD_{600}$ of 0.3. These cultures were incubated at 30°C with shaking for 2-3 hours. This culture was then diluted and plated on YPD and then replica plated onto SC-5FOA (25% w/v g Bacto-Agar, 6.72% w/v YNB, 1% v/v mL 20x KS supplement without URA, 2% v/v glucose, 10 mL 5-Fluoroorotic Acid (Zymo Research), 50 mg uracil (MP Biomedicals 103204). 5FOA resistant colonies were checked for excision of the KlURA3 marker using colony PCR. Transformants with klURA3 excised were grown in liquid YPD to saturation twice and then plated on YPD for ~100 colonies per plate. These were replica plated onto YPD + 10µg/ml phleomycin. Phleomycin sensitive colonies (colonies that had lost the plasmid pMM296) were reconfirmed by colony PCR to have loxed out klURA3. This generated yMM1472 (Matα trp1Δ63 leu2Δ1 ura3::pRPL18B-CIB1VP16-tENO1-pRPL18B-ZCRY2PHR-pZF(3BS)-mRuby2-loxPScar).

To test that neither the loxP-KlURA3-loxP marker nor removal of the klURA3 affected the function of ZCRY2PHR or CIB1VP16 we compared mRuby2 induction in yMM1462, yMM1468, and yMM1472. These strains were grown overnight in low fluorescence media and then diluted back to $OD_{600}$ 0.01 in the LPA in the morning. Cultures were run in triplicate in the LPA at 15µW blue-light overnight (16 hours). Fluorescence was assessed by flow cytometry. As seen in **Supplemental Figure 9,** all three strains induce at comparable levels.

### References

[1] R. Gietz and R. Schiestl, "High-efficiency yeast transformation using the LiAc/SS carrier DNA/PEG method," *Nature Protocols,* vol. 2, pp. 31-34, 2007.

[2] S. Harju, H. Fedosyuk and K. Peterson, "Rapid isolation of yeast genomic DNA: Bust n' Grab," *BMC Biotechnology,* vol. 4, no. 8, 2004.

[3] D. Amberg, D. Burke and J. Strathern, Methods in yeast genetics : a Cold Spring Harbor Laboratory course manual, Cold Spring Harbor, N.Y. : Cold Spring Harbor Laboratory Press, 2005.

[4] J. Sambrook and D. Russell, "The Inoue Method for Preparation and Transformation of Competent E. Coli: "Ultra-competent" cells," *CSH Protocols,* no. 1, 2006.

[5] K. Gerhardt, E. Olson, S. Castillo-Hair, L. Hartsough, B. Landry, F. Ekness, R. Yokoo, E. Gomez, P. Ramakrishnan, J. Suh, D. Savage and J. Tabor, "An open-hardware platform for optogenetics and photobiology," *Scientific Reports,* vol. 6, no. 35363, 2016.

[6] K. Sweeney, N. Moreno Morales, Z. Burmeister, A. Numunkar and M. McClean, "Easy calibration of the Light Plate Appartus for optogenetic experiments," *MethodsX,* 2019.

[7] E. Andersen, "tPCR-Directed In Vivo Plasmid Construction Using Homologous Recombination in Baker’s Yeast," *Molecular Methods for Evolutionary Genetics. Methods in Molecular Biology ,* vol. 772, pp. 409-421, 8 7 2011.

[8] R. McIsaac, S. Silverman, M. McClean, P. Gibney, J. Macinskas, M. Hickman, A. Petti and a. D. Botstein, "Fast-acting and nearly gratuitous induction of gene expression and protein depletion in *Saccharomyces cerevisiae*.," *Molecular Biology of the Cell,* vol. 22, pp. 4447-4459, 2011.

[9] R. McIsaac, B. Oakes, X. Wang, K. Dummit, D. Botstein and M. Noyes, "Synthetic gene expression perturbation systems with rapid, tunable single-gene specificity in yeast," *Nucleic Acids Research,* vol. 23, pp. 2993-3008, 2012.

[10] R. Sikorski and P. Heiter, "A system of shuttle vectors and yeast host strains designed for efficient manipulation of DNA in *Saccharomyces cerevisiae*," *Genetics,* vol. 122, no. 1, pp. 19-27, 1989.

[11] M. Sheff and K. Thorn, "Optimized cassettes for fluorescent protein tagging in *Saccharomyces cerevisiae*," *Yeast,* vol. 21, pp. 661-670, 2004.

[12] M. E. Lee, W. Deloache, D. Cervantes and J. Dueber, "A highly characterized yeast toolkit for modular, multipart assembly," *ACS Synthetic Biology,* vol. 4, no. 9, pp. 975-986, 2015.

[13] M. Zwietering, I. Jongenburger, F. Rombouts and K. Van't Riet, "Modeling the bacterial growth curve," *Applied and environmental microbiology,* vol. 56, no. 6, pp. 1875-1881, 1990.

[14] E. Jones, E. Oliphant, P. Peterson and e. al, "SciPy: Open Source Scientific Tools for Python," 2001. [Online]. Available: http://www.scipy.org/. [Accessed 26 2 2019].

[15] M. Branch, T. Coleman and Y. Li, "A Subspace, Interior, and Conjugate Gradient Method for Large-Scale Bound-Constrained Minimization Problems," *SIAM Journal of Scientific Computing,* vol. 21, no. 1, pp. 1-23, 1999.

[16] Z. Carter and D. Delneri, "New generation of loxP-mutated deletion cassettes for the genetic manipulation of yeast natural isolates," *Yeast,* vol. 27, pp. 765-775, 2010.

[17] C. Brachmann, A. Davies, G. Cost, E. Caputo, J. Li, P. Heiter and J. Boeke, "Designer deletion strains derived from *Saccharomyces cerevisiae* S288C: a useful set of strains and plasmids for PCR-mediated gene disruption and other applications," *Yeast,* vol. 14, pp. 115-132, 1998.

[18] U. Gueldener, J. Heinisch, G. Koehler, D. Voss and J. Hegemann, "A second set of loxP marker cassettes for Cre-mediated multiple gene knockouts in budding yeast," *Nucleic acids research,* vol. 30, no. 6, 2002.

### Supplemental Figures

#### Supplemental Figure Captions

**Supplemental Figure 1 pZF Promoter Sequences**: Sequences of the $P_{GAL1}$ promoter compared with the synthetic promoters containing variable numbers and orientations of binding sites for the Zif268DBD.

**Supplemental Figure 2 pZF Promoters:** Comparison of the synthetic promoters. (a) Strain yMM1146 (Mat α trp1Δ63 leu2Δ1 ura3-52) was co-transformed with pGAL4AD-CIB1 (pMM159), pZDBD-CRY2 (pMM284) and one of the reporter plasmids (pMM285-290) with promoters as detailed in **Supplemental Fig. 1**. Cells were induced with blue-light for up to 17 hours. Expression of yEVENUS was measured using flow cytometry. (**b**) Strain yMM1146 was transformed with the reporter plasmid as well as pMM315 and pMM316, which contain the appropriate auxotrophic markers, but no CRY2 or CIB1 constructs. These strains were induced with blue-light as in (**a**). The synthetic promoters alone in the presence of blue-light do not lead to yEVENUS induction.

**Supplemental Figure 3 Crosstalk:** Strain yMM1146 was transformed with the indicated constructs and exposed to blue-light for 17 hours as described in the **Methods**. Expression of yEVENUS was measured using flow cytometry. Empty vector (ø) controls for DBD-CRY2, AD-CIB1, and the promoter plasmids are respectively pMM315 (pGAL4BD TRP1 CEN), pMM316 (pGAL4AD LEU2 CEN), and pMM8 (URA3 CEN).

**Supplemental Figure 4 Linkers**: Quantification of the effect of linker length between Zif268DBD and the CRY2 protein (**a**) Strain yMM1146 (Mat alpha trp1Δ63 leu2Δ1 ura3-52) was co-transformed with the pGAL4AD-CIB1 plasmid (pMM159), the p4BS*-yEVENUS plasmid (pMM289) and pZDBD-CRY2 with three different linker lengths (pMM282, pMM283, pMM284). Cells were grown to mid-log in SC-Ura-Leu-Trp before being exposed to blue-light for 17 hours as detailed in the **Methods** (**b**) Details of the linkers used between the Zif268 DNA-binding domain (ZDBD) and CRY2.

**Supplemental Figure 5 Nuclear Localization Signal:** We used two different nuclear localization signal (NLS) sequences to localize the Z-CRY2 construct to the nucleus. The SV40 sequence (P K K K R K V) was taken from pMM159 (GAL4AD-CIB1), while the H2B nuclear localization sequence (G K K R S K A K) is based on the *Saccharomyces cerevisiae* histone

H2B sequence. Cells were transformed with the p4BS*-yEVENUS reporter plasmid (pMM289), pGAL4AD-CIB1 (pMM159) or pVP16-CIB1 (pMM281), and SV40NLS-Z-CRY2 (pMM313) or H2BNLS-Z-CRY2 (pMM314) and exposed to blue-light for up to 17 hours. The induction of yEVENUS induction was measured using flow cytometry, as detailed in the **Methods.**

**Supplemental Figure 6 Yeast Optogenetic Toolkit Parts: (a)** Optogenetic parts created in this paper that integrate with the Lee, *et al* Yeast Toolkit [12] are highlighted and shown with additional Yeast Toolkit parts used in this paper. (**b**) A preassembled entry vector for integration at the LEU2 locus. A transcriptional unit (promoter, coding sequence, terminator) can be assembled into this vector replacing the BsaI-flanked GFP dropout. (**c**) A similar preassembled integration vector but using the loxP-KlURA3-loxP yeast selection marker. Once integrated, the KlURA3 selection marker can be removed by CRE-recombinase expression as detailed in the **Methods**.

**Supplemental Figure 7 Dosage Strains: (A)** Comparison of pZF-mRUBY2 expression in yeast strains containing ZDBD-CRY2 and VP16-CIB1 expressed under different strength (High-pTEF1, Medium-pRPL18B, Low-pRNR2) yeast promoters compared to strains with constitutive expression of mRUBY2 (S-pTDH3, M-pRPL18B, W-pREV1). This is the raw fluorescence data that corresponds to the fold-change data displayed in **Figure 2D**. **(B)** Endpoint $OD_{600}$ in the strains shown in **(A)**. (**C**) Growth rate, μ, of the strains shown in (**A**).

**Supplemental Figure 8 Timecourse**: Timecourse of gene expression in the ZDBD-$CRY2_{medium}$/VP16-$CIB1_{medium}$ strain (**A,C**) and the ZDBD-$CRY2_{medium}$/VP16-$CIB1_{high}$ (**B,D**) strain compared in dark (black) and 15μW/$cm^2$ light (red). Time is in hours after turning on the light in the Light Plate Apparatus. Two separate timecourses show early induction (0-6 hours, **A,B**) as well as the approach to saturation (6-20 hours **C, D**).

**Supplemental Figure 9 Marker Recycling:** Integration of the ZDBD-$CRY2_{medium}$/VP16-$CIB1_{medium}$ pZF(3BS)-mRUBY2 cassette using **(A)** pMM641 (scURA3 selection marker) and **(B)** pMM647 (klURA3 selection marker) were compared for their ability to induce mRUBY2 expression in response to blue light. The choice of marker (yMM1462/scURA3 vs yMM1468/klURA3) or recycling of the klURA3 marker using CRE-recombinase expression (yMM1472) did not affect the optogenetic system as indicated by comparable mRuby2 fluorescence after growing the different strains in blue light (or dark) for 16 hours. Means of the induced and uninduced populations were determined by fitting the flow cytometry data to a lognormal distribution. Error bars represent S.E.M.

#### Supplemental Tables

Additional details on yeast strains, oligos, and plasmids can be found in the **Supplemental Material.**

##### Supplemental Table 1: Yeast Strains

| ID | Alias | Genotype | Description | Source/ Reference |
|---|---|---|---|---|
| **yMM83** | ySR32, BY4741 | MATa his3Δ1 leu2Δ0 met15Δ0 ura3Δ0 | Assembly strain | Brachmann, CB et al (1998) [17] |
| **yMM1008** | pGAL1-ARO80 | (PGAL10+gal1)Δ::loxP gal4Δ::LEU2 HAP1 leu2Δ0::PACT1-GEV-NatMX KANMX-PGAL1-ARO80 | Source of VP16 from GEV | McClean Lab |
| **yMM1146** | DBY8750, KSY1284 | Mat alpha trp1Δ63 leu2Δ1 ura3-52 | Assembly and integration strain | Botstein lab |
| **yMM1367** | yMM1146_ pMM364@ HO | Mat alpha trp1Δ63 leu2Δ1 ura3-52 HO::SV40NLS-VP16-CIB1 loxP-KIURA3-loxP SV40NLS-Zif268DBD-CRY2PHR | Strain with integrated optogenetic system, to allow for induction of GOI using KanMXREV-pZF promoter | This study |

| | | | | |
|---|---|---|---|---|
| **yMM1401** | mRuby2 HIGH | Mat alpha trp1Δ63 leu2Δ1::LEU2-pTDH3-mRUBY2-tADH1 ura3-52 HO::S40NLS-VP16-CIB1 loxP-KIURA3-loxP SV40NLS-Zif268DBD-CRY2PHR | Benchmark for high mRuby2 expression (pTDH3) | This study |
| **yMM1403** | mRuby2 LOW | Mat alpha trp1Δ63 leu2Δ1::LEU2-pREV1-mRUBY2-tADH1 ura3-52 HO::S40NLS-VP16-CIB1 loxP-KIURA3-loxP SV40NLS-Zif268DBD-CRY2PHR | Benchmark for low mRuby2 expression (pREV1) | This study |
| **yMM1444** | mRuby2 MEDIUM | Mat alpha trp1Δ63 leu2Δ1::LEU2-pRPL18B-mRUBY2-tADH1 ura3-52 HO::S40NLS-VP16-CIB1 loxP-KIURA3-loxP SV40NLS-Zif268DBD-CRY2PHR | Benchmark for medium mRuby2 expression (pRPL18B) | This Study |
| **yMM1458** | yMA001 | Matα trp1Δ63 leu2Δ1 ura3-52::pTEF1-SV40NLS-VP16_CIB1-tENO1-Scar1-pTEF1-SV40NLS-ZifDBDCRY2PHR-tSSA1-Scar2-pZF(3BS)-mRuby2-tADH1-ScarRE-scURA3 | High/HighCIB1AD/ ZCRY2 expressing strain | This Study |
| **yMM1459** | yMA002 | Matα trp1Δ63 leu2Δ1 ura3-52::pTEF1-SV40NLS-VP16_CIB1-tENO1-Scar1-pRPL18B-SV40NLS-ZifDBDCRY2PHR-tSSA1-Scar2-pZF(3BS)-mRuby2-tADH1-ScarRE-scURA3 | High/Medium CIB1AD/ZCRY2 expressing strain | This Study |
| **yMM1460** | yMA003 | Matα trp1Δ63 leu2Δ1 ura3-52::pTEF1-SV40NLS-VP16_CIB1-tENO1-Scar1-pRNR2-SV40NLS-ZifDBDCRY2PHR-tSSA1-Scar2-pZF(3BS)-mRuby2-tADH1-ScarRE-scURA3 | High/Low CIB1AD/ZCRY2 expressing strain | This Study |
| **yMM1461** | yMA004 | Matα trp1Δ63 leu2Δ1 ura3-52::pRPL18B-SV40NLS-VP16_CIB1-tENO1-Scar1-pTEF1-SV40NLS-ZifDBDCRY2PHR-tSSA1-Scar2-pZF(3BS)-mRuby2-tADH1-ScarRE-scURA3 | Medium/High CIB1AD/ZCRY2 expressing strain | This Study |
| **yMM1462** | yMA005 | Matα trp1Δ63 leu2Δ1 ura3-52::pRPL18B-SV40NLS-VP16_CIB1-tENO1-Scar1-pRPL18B-SV40NLS-ZifDBDCRY2PHR-tSSA1-Scar2-pZF(3BS)-mRuby2-tADH1-ScarRE-scURA3 | Medium/Medium CIB1AD/ZCRY2 expressing strain | This Study |
| **yMM1463** | yMA006 | Matα trp1Δ63 leu2Δ1 ura3-52::pRPL18B-SV40NLS-VP16_CIB1-tENO1-Scar1-pRNR2-SV40NLS-ZifDBDCRY2PHR-tSSA1-Scar2-pZF(3BS)-mRuby2-tADH1-ScarRE-scURA3 | Medium/Low CIB1AD/ZCRY2 expressing strain | This Study |
| **yMM1464** | yMA007 | Matα trp1Δ63 leu2Δ1 ura3-52::pRNR2-SV40NLS-VP16_CIB1-tENO1-Scar1-pTEF1-SV40NLS-ZifDBDCRY2PHR-tSSA1-Scar2-pZF(3BS)-mRuby2-tADH1-ScarRE-scURA3 | Low/High CIB1AD/ ZCRY2 expressing strain | This Study |
| **yMM1465** | yMA008 | Matα trp1Δ63 leu2Δ1 ura3-52::pRNR2-SV40NLS-VP16_CIB1-tENO1-Scar1-pRPL18B-SV40NLS-ZifDBDCRY2PHR-tSSA1-Scar2-pZF(3BS)-mRuby2-tADH1-ScarRE-scURA3 | Low/Medium CIB1AD/ZCRY2 expressing strain | This Study |
| **yMM1466** | yMA009 | Matα trp1Δ63 leu2Δ1 ura3-52::pRNR2-SV40NLS-VP16_CIB1-tENO1-Scar1-pRNR2-SV40NLS-ZifDBDCRY2PHR-tSSA1-Scar2-pZF(3BS)-mRuby2-tADH1-ScarRE-scURA3 | Low/Low CIB1AD/ZCRY2 expressing strain | This Study |
| **yMM1467** | yMA011 | Matα trp1Δ63 leu2Δ1 ura3-52::pTEF1-SV40NLS-VP16_CIB1-tENO1-Scar1-pRPL18B-SV40NLS-ZifDBDCRY2PHR-tSSA1-Scar2-pZF(3BS)-mRuby2-tADH1-ScarRE-loxP-KLURA3-loxP | High/Medium CIB1AD/ZCRY2 expressing strain w/loxP-klURA3 | This Study |
| **yMM1468** | yMA014 | Matα trp1Δ63 leu2Δ1 ura3-52::pRPL18B-SV40NLS-VP16_CIB1-tENO1-Scar1-pRPL18B-SV40NLS-ZifDBDCRY2PHR-tSSA1-Scar2-pZF(3BS)-mRuby2-tADH1-ScarRE-loxP-KLURA3-loxP | Medium/Medium CIB1AD/ZCRY2 expressing strain | This Study |
| **yMM1472** | yMA032 | Matα trp1Δ63 leu2Δ1 ura3-52::pRPL18B-SV40NLS-VP16_CIB1-tENO1-Scar1-pRPL18B-SV40NLS- | Medium/Medium CIB1AD/ZCRY2 expressing strain | This Study |

| | | | | |
|---|---|---|---|---|
| | | ZifDBDCRY2PHR-tSSA1-Scar2-pZF(3BS)-mRuby2-tADH1-ScarRE-loxPScar | | |
| **yMM1473** | yMA037 | Matα trp1Δ63 leu2Δ1 ura3-52::pTEF1-SV40NLS-VP16_CIB1-tENO1-Scar1-pRPL18B-SV40NLS-ZifDBDCRY2PHR-tSSA1-Scar2-spacer-ScarRE-scURA3 | No fluorescence control for yMM1458 | This Study |
| **yMM1474** | yMA038 | Matα trp1Δ63 leu2Δ1 ura3-52::pTEF1-SV40NLS-VP16_CIB1-tENO1-Scar1-pRPL18B-SV40NLS-ZifDBDCRY2PHR-tSSA1-Scar2-pREV1-mRUBY2-tADH1-ScarRE-scURA3 | Low mRuby2 fluorescence control for yMM1458 | This Study |
| **yMM1475** | yMA039 | Matα trp1Δ63 leu2Δ1 ura3-52::pTEF1-SV40NLS-VP16_CIB1-tENO1-Scar1-pRPL18B-SV40NLS-ZifDBDCRY2PHR-tSSA1-Scar2-pRPL18B-mRUBY2-tADH1-ScarRE-scURA3 | Medium mRuby2 fluorescence control for yMM1458 | This Study |
| **yMM1476** | yMA040 | Matα trp1Δ63 leu2Δ1 ura3-52::pTEF1-SV40NLS-VP16_CIB1-tENO1-Scar1-pRPL18B-SV40NLS-ZifDBDCRY2PHR-tSSA1-Scar2-pTDH3-mRUBY2-tADH1-ScarRE-scURA3 | High mRuby2 fluorescence control for yMM1458 | This Study |
| **yMM1477** | yMA041 | Matα trp1Δ63 leu2Δ1 ura3-52::pRPL18B-SV40NLS-VP16_CIB1-tENO1-Scar1-pRPL18B-SV40NLS-ZifDBDCRY2PHR-tSSA1-Scar2-Spacer-ScarRE-scURA3 | No fluorescence control for yMM1459 | This Study |
| **yMM1478** | yMA042 | Matα trp1Δ63 leu2Δ1 ura3-52::pRPL18B-SV40NLS-VP16_CIB1-tENO1-Scar1-pRPL18B-SV40NLS-ZifDBDCRY2PHR-tSSA1-Scar2-pREV1-mRUBY2-tADH1-ScarRE-scURA3 | Low mRuby2 fluorescence control for yMM1459 | This Study |
| **yMM1479** | yMA043 | Matα trp1Δ63 leu2Δ1 ura3-52::pRPL18B-SV40NLS-VP16_CIB1-tENO1-Scar1-pRPL18B-SV40NLS-ZifDBDCRY2PHR-tSSA1-Scar2-pRPL18B-mRUBY2-tADH1-ScarRE-scURA3 | Medium mRuby2 fluorescence control for yMM1459 | This Study |
| **yMM1480** | yMA044 | Matα trp1Δ63 leu2Δ1 ura3-52::pRPL18B-SV40NLS-VP16_CIB1-tENO1-Scar1-pRPL18B-SV40NLS-ZifDBDCRY2PHR-tSSA1-Scar2-pTDH3-mRUBY2-tADH1-ScarRE-scURA3 | High mRuby2 fluorescence control for yMM1459 | This Study |

#### Supplemental Table 2: Primer used in this study

| oMM | Alias | Sequence | Target | Purpose (Brief) |
|---|---|---|---|---|
| **oMM0166** | rev_pGal4AD-CIB1_check | aagtgaacttgcggggtttt | CIB1, CIB1N | Amplification/Sequencing |
| **oMM0186** | tADH1_rev | CCGGTAGAGGTGTGGTCAAT | tADH1 | Sequencing |
| **oMM0212** | Forward_pGAL1 | GTTTTCCCAGTCACGACGTT | pGAL1 | Sequencing |
| **oMM0214** | Forward_pGAL1 | ACGCTTAACTGCTCATTGCT | pGAL1 | Sequencing |
| **oMM0250** | Lox_General_for | cgtacgctgcaggtcgac | loxP_KIURA3-loxP | Amplification |
| **oMM0251** | Lox_General_rev | cactatagggagaccggcag | loxP_KIURA3-loxP | Amplification |
| **oMM0323** | pRS415_for | cccagtcacgacgttgtaaaacg | pRS415 | Sequencing |
| **oMM0349** | URA3_reverse | URA3 | CCCTACACGTTCGCTATGCT | Sequencing |
| **oMM0354** | pADH1_forward | TCGTTGTTCCAGAGCTGATG | pADH1 | Sequencing |
| **oMM0383** | CRY2_ZFBD_tem2_for | tgcggccgcaatacaatagctagcttcatgaagatggacaaaa | CRY2-tADH1 | Cloning |
| **oMM0384** | CRY2_ZFBD_tem2_rev | ataagagctccatgccggtagaggtgt | CRY2-tADH1 | Cloning |

| | | | | |
|---|---|---|---|---|
| **oMM0393** | Seq_3 | GTGAGCAGCTGCTGCTAATG | CRY2 | Sequencing |
| **oMM0399** | For_Internal_CRY2 | CAGTCTTGCTCGTTGGCATC | CRY2 | Sequencing |
| **oMM0400** | For_GAL4AD_VP16_swap | tccaaaaaagaagagaaaggtcgaattgggtaccgccgccTCGGAGCTC CACTTAGACGG | GAL4AD,VP16 | Yeast Recombinational Cloning |
| **oMM0401** | Rev_GAL4AD_VP16_swap | cgctagcttcggcgctcgccctatagtgagtcgtattaaaCCCACCGTACTC GTCAATTC | GAL4AD,VP16 | Yeast Recombinational Cloning |
| **oMM0402** | GAL4AD_forward | GCGTATAACGCGTTTGGAAT | GAL4AD | Colony PCR, Sequencing |
| **oMM0403** | CIB1_reverse | CGGAAAGAACCATTTCAGGA | CIB1 | Colony PCR, Sequencing |
| **oMM0404** | VP16_forward | TTTCGATCTGGACATGTTGG | VP16 | Colony PCR, Sequencing |
| **oMM0405** | pADH1_seq | TTCCTTCATTCACGCACACT | pyADH1 | Sequencing |
| **oMM0407** | Rev_ZFBD-CRY2 linker3 | ataagctagcACCTGTATGGATTTTGGTATG | FLAG(3X)-NLS-Zif268DBD | Cloning |
| **oMM0408** | Rev_ZFBD-CRY2 linker4pro | ataagctagctggACCTGTATGGATTTTGGTATG | FLAG(3X)-NLS-Zif268DBD | Cloning |
| **oMM0409** | Rev_ZFBD-CRY2 linker10 | ataagctagctccgccaccactgccacctccACCTGTATGGATTTTGGT ATG | FLAG(3X)-NLS-Zif268DBD | Cloning |
| **oMM0413** | ZIF1_Top | ggccgcGCGTGGGCGt | Zif268 Binding Sites | Cloning of ZF BS |
| **oMM0414** | ZIF1_Bottom | ctagaCGCCCACGCgc | Zif268 Binding Sites | Cloning of ZF BS |
| **oMM0415** | Zif3_Top | ggccgcGCGTGGGCGAGCGTGGGCGAGCGTGGGCGt | Zif268 Binding Sites | Cloning of ZF BS |
| **oMM0416** | Zif3_Bottom | ctagaCGCCCACGCTCGCCCACGCTCGCCCACGCgc | Zif268 Binding Sites | Cloning of ZF BS |
| **oMM0417** | Zif3_overlap2_top | ggccgcGCGTGGGCGTGGGCGGCGTGGGCGt | Zif268 Binding Sites | Cloning of ZF BS |
| **oMM0418** | Zif3_overlap2_bottom | ctagaCGCCCACGCCGCCCACGCCCACGCgc | Zif268 Binding Sites | Cloning of ZF BS |
| **oMM0419** | Zif3_invert2_top | ggccgcGCGTGGGCGAGCGGGTGCGt | Zif268 Binding Sites | Cloning of ZF BS |
| **oMM0420** | Zif3_intert2_bottom | ctagaCGCACCCGCTCGCCCACGCgc | Zif268 Binding Sites | Cloning of ZF BS |
| **oMM0421** | XmaI-5'-Venus | ttatcccgggatgtctaaaggtgaagaattat | Venus | Cloning |
| **oMM0423** | AscI-3'-Venus | ataaggcgcgccCTAttatttgtacaattcatccatac | Venus | Cloning |
| **oMM0456** | ZCRY ADH 5p (NgoMIV) | ttatgccggcCAACTTCTTTTCTTTTTTTTTCT | pADH1 | Cloning |
| **oMM0457** | ZCRY ADH3p (spacer-not1) | gcggccgcaggcttgcttcaagcttGGAGTTGATTGTATGCTTGG | pADH1 | Cloning |
| **oMM0458** | NLS 5p (ADH spacer homology-NotI) | caagcctgcggccgcATGGATTACAAGGATGACGA | FLAG(3X)-NLS-Zif268DBD | Cloning |
| **oMM0481** | Zif 3corrected_top | ggccgcGCGTGGGCGTGCGTGGGCGTGCGTGGGCGt | Zif268 Binding Sites | Cloning of ZF BS |
| **oMM0482** | Zif 3corrected_bottom | ctagaCGCCCACGCACGCCCACGCACGCCCACGCgc | Zif268 Binding Sites | Cloning of ZF BS |
| **oMM0483** | Zif 4_top | ggccgcGCGTGGGCGTGCGTGGGCGTGCGTGGGCGTGCGTG GGCGt | Zif268 Binding Sites | Cloning of ZF BS |
| **oMM0484** | Zif 4_bottom | ctagaCGCCCACGCACGCCCACGCACGCCCACGCACGCCCAC GCgc | Zif268 Binding Sites | Cloning of ZF BS |
| **oMM0485** | Zif opp3_top | ggccgcCGCCCACGCACGCCCACGCACGCCCACGCt | Zif268 Binding Sites | Cloning of ZF BS |
| **oMM0486** | Zif opp3_bottom | ctagaGCGTGGGCGTGCGTGGGCGTGCGTGGGCGgc | Zif268 Binding Sites | Cloning of ZF BS |
| **oMM0535** | ZCRY2_promswap_noAT Gmut_for | CTGCACAATATTTCAAGCTATACCAAGCATACAATCAACTAT CTCATATACAATGGATTA | ZDBD-CRY2 | Yeast Recombinational Cloning |

| | | | | |
|---|---|---|---|---|
| **oMM0536** | ZCRY2_promswap_noATGmut_rev | ATCGTCCTTATAGTCCCCGGTCTTATCGTCGTCATCCTTGTAATCCATTGTATATGAGAT | ZDBD-CRY2 | Yeast Recombinational Cloning |
| **oMM0538** | ZCRY2_promswap_insertPACI_for | CTGCACAATATTTCAAGCTATACCAAGCATACAATCAACTttaattaaATGGATTACAAG | ZDBD-CRY2 | Yeast Recombinational Cloning |
| **oMM0539** | ZCRY2_promswap_insertPACI_rev | CATCATCGTCCTTATAGTCCCCGGTCTTATCGTCGTCATCCTTGTAATCCATTTAATTAA | ZDBD-CRY2 | Yeast Recombinational Cloning |
| **oMM0547** | pMM159_exCIB1_for | AGATCTTTAATACGACTCACTATAGGGCGAcatcgagctcgagctgcagatgaatcgtag | CIB1 | Yeast Recombinational Cloning |
| **oMM0548** | pMM159_exCIB1_rev | CTACGATTCATCTGCAGCTCGAGCTCGATGTCGCCCTATAGTGAGTCGTATTAAAGATCT | CIB1 | Yeast Recombinational Cloning |
| **oMM0549** | pMM160_exCRY2_for | AACAAAGGTCAAAGACAGTTGACTGTATCGggcaagtgcacaaacaatacttaaataaat | CRY2 | Yeast Recombinational Cloning |
| **oMM0550** | pMM160_exCRY2_rev | ATTTATTTAAGTATTGTTTGTGCACTTGCCCGATACAGTCAACTGTCTTTGACCTTTGTT | CRY2 | Yeast Recombinational Cloning |
| **oMM0551** | SV40NLS_YRC_Forward | CTGCACAATATTTCAAGCTATACCAAGCATACAATCAACTATCTCATATACAATGGCCCC | Zif268DBD-CRY2 | Yeast Recombinational Cloning |
| **oMM0552** | SV40NLS_YRC_Reverse | CCCCGTGAATACCAACCTTCCTCTTCTTCTTGGGGGCCATTGTATATGAGAT | Zif268DBD-CRY2 | Yeast Recombinational Cloning |
| **oMM0553** | SV40NLS_cterm_YRC_Forward | TTACTACAAGTTTGGGAAAAAATGGTTGCAAAGCCCCCAAGAAGAAGAGGAAGGTTGTGA | CRY2, SV40NLS | Yeast Recombinational Cloning |
| **oMM0554** | SV40NLS_cterm_YRC_Reverse | TTATTTAATAATAAAAATCATAAATCATAAGAAATTCGCCTCACAACCTTCCTCTTCTTC | CRY2, SV40NLS | Yeast Recombinational Cloning |
| **oMM0562** | CRY2->CRY2PHR_forward_YRC | TTCAAGAACCCGTGAAGCACAGATCATGATCGGAGCAGCAgcccgggtcgacctgcagcc | CRY2, CRY2PHR | Yeast Recombinational Cloning |
| **oMM0563** | CRY2->CRY2PHR_reverse_YRC | AAAAATCATAAATCATAAGAAATTCGCCCGGAATTAGCTTGGCTGCAGGTCGACCCGGGC | CRY2, CRY2PHR | Yeast Recombinational Cloning |
| **oMM0564** | CRY2->CRY2PHR-SV40NLS_forward_YRC | GAACCCGTGAAGCACAGATCATGATCGGAGCAGCAGCCCCCAAGAAGAAGAGGAAGGTTG | CRY2, CRY2PHR | Yeast Recombinational Cloning |
| **oMM0565** | CRY2->CRY2PHR-SV40NLS_reverse_YRC | AAAAATCATAAATCATAAGAAATTCGCCCGGAATTAGCTTCAACCTTCCTCTTCTTCTTG | CRY2, CRY2PHR | Yeast Recombinational Cloning |
| **oMM0566** | SV40_NoFlag_NotI_KpnI_forward | ggccgcATGGCtCCCAAGAAGAAGAGGAAGGTTGGTATTCACGGGggtac | SV40 | Cloning |
| **oMM0567** | SV40_NoFlag_NotI_KpnI_reverse | cCCCGTGAATACCAACCTTCCTCTTCTTCTTGGGaGCCATgc | SV40 | Cloning |
| **oMM0568** | Histone 2B NLS w/GlyPro Linker 4x_NotI_KpnI_forward | ggccgcATGGGTAAGAAGAGATCTAAGGCTAAGGGACCAGGTCCTGGACCAGGACCTGGCGGAGGCggtac | H2B NLS | Cloning |
| **oMM0569** | Histone 2B NLS w/GlyPro Linker 4x_NotI_KpnI_reverse | cGCCTCCGCCAGGTCCTGGTCCAGGACCTGGTCCCTTAGCCTTAGATCTCTTCTTACCCATgc | H2B NLS | Cloning |
| **oMM0684** | KLURA3_rev | KLURA3 | GAATCAGCGCTCCCCATTAA | Sequencing |
| **oMM0776** | pZF Part Forward | GCATCGTCTCATCGGTCTCAAACGcgaggcaagctaaacagat | pZF | Cloning |
| **oMM0870** | Gene_plasmid_for | CGGATGACACGAACTCACGA | ConE | Sequencing |
| **oMM0871** | Gene_plasmid_rev | GGTTCCGGCTGTCTTGCTTA | ConS | Sequencing |

| | | | | |
|---|---|---|---|---|
| **oMM0991** | loxP-KLURA3_for | GCATCGTCTCATCGGTCTCATACAGCAGGTCGACAACCCTTA | loxP-KIURA3-loxP | Cloning |
| **oMM0992** | loxP-KLURA3_rev | ATGCCGTCTCAGGTCTCAACTCAGTGGATCTGATATCACCTA ATAA | loxP-KIURA3-loxP | Cloning |
| **oMM1003** | SV40NLS'_ZiF_CRY2PHR forward | GCATCGTCTCATCGGTCTCATATGgcccccaagaagaagaggaa | SV40NLS-Zif268DBD-CRY2PHR | Cloning |
| **oMM1039** | pZiF_Rev | ATGCCGTCTCAGGTCTCACATAgtacctatagttttttctccttgac | pZF | Cloning |
| **oMM1065** | pZiF_F | TTGAAGTGCGcgCGCGCGTGGG | pZF(3BS) | Site-directed mutagenesis |
| **oMM1066** | pZiF_R | TATAGCAATGAGCAGTTAAGCG | pZF(3BS) | Site-directed mutagenesis |
| **oMM1067** | pCIB1_F | TGTACGGTGActCGACGGTGGAAG | CIB1 | Site-directed mutagenesis |
| **oMM1068** | pCIB1_R | TCATCGGAAGATTCAAAC | CIB1 | Site-directed mutagenesis |
| **oMM1069** | pCRY2_1_F | GTTTAGAAGActCCTAAGGATTGAGGATAATC | CRY2PHR | Site-directed mutagenesis |
| **oMM1070** | pCRY2_1_R | CAAACTATAGTCTTTTTGTCC | CRY2PHR | Site-directed mutagenesis |
| **oMM1071** | pCRY2_2_F | TAGAGCTTGGagTCCAGGATGG | CRY2PHR | Site-directed mutagenesis |
| **oMM1072** | pCRY2_2_R | GTTAACAACGCATTGCTC | CRY2PHR | Site-directed mutagenesis |
| **oMM1073** | CRY2_REV | ATGCCGTCTCAGGTCTCAGGATCCtttgcaaccattttttcccaaac | CRY2 | Cloning |
| **oMM1074** | CRY2PHR_REV | ATGCCGTCTCAGGTCTCAGGATCCCCGGGCtgctgctccgatcatg at | SV40NLS-Zif268DBD-CRY2PHR | Cloning |
| **oMM1124** | pMM281_mutate_for | TGTACGGTGAaACGACGGTGG | CIB1 | Site-directed mutagenesis |
| **oMM1125** | pMM281_mutate_rev | TCATCGGAAGATTCAAACCGGC | CIB1 | Site-directed mutagenesis |
| **oMM1126** | pMM515_mutate_for | GTTTAGAAGAgatCTAAGGATTGAGGATAATCC | CRY2 | Site-directed mutagenesis |
| **oMM1127** | pMM515_mutate_rev | CAAACTATAGTCTTTTTGTCC | CRY2 | Site-directed mutagenesis |
| **oMM1137** | pMM491_rev | TCACAGGCTTAGGTGGATCT | ConR1 | Sequencing |
| **oMM1138** | pMM532_Rev | CATGATGACCGCACTGACTG | ConL1 | Sequencing |
| **oMM1141** | pMM537_rev | CGGATCACCGTACTATGTGTGA | ConR2 | Sequencing |
| **oMM1142** | pMM533_fwd | ACAGGAGCAAGCGCGATAGG | ConL2 | Sequencing |
| **oMM1146** | pMM534_fwd | ATGCCGATGCACGCTCATAT | ConL3 | Sequencing |
| **oMM1149** | pMM539_rev | GTTAGCCTGCCTCGATTTCA | ConR4 | Sequencing |
| **oMM1150** | pMM535_fwd | GATGACGTAACACCGAGCCA | ConL4 | Sequencing |
| **oMM1174** | tENO1_for | cgcgtgtatccgcccgc | tENO1 | Sequencing |
| **oMM1175** | tSSA1_for | gccaattggtgcggcaattg | tSSA1 | Sequencing |
| **oMM1176** | scURA3_rev | ggactaggatgagtagcagcacgttcc | scURA3 | Sequencing |
| **oMM1177** | scURA3_5' | gtggctgtggtttcagggtccat | scURA3 | Sequencing |
| **oMM1220** | tADH1_forward | TGCAAATCGCTCCCCATT | tADH1 | Sequencing |

#### Supplemental Table 3: Plasmids used in this study

| ID | Alias | Gene(s) or Insert Name | Yeast Marker | Bacterial Resistance | Source/ Reference |
|---|---|---|---|---|---|

| | | | | | |
|---|---|---|---|---|---|
| **pMM0006** | pRS414 | TRP1 CEN6 ARS4 | TRP1 | Ampicillin | Sikorski and Heiter, 1989 |
| **pMM0008** | pRS416 | URA3 CEN6 ARS4 | URA3 | Ampicillin | Sikorski and Heiter, 1989 |
| **pMM0159** | pGal4AD-CIB1 | pscADH1-GAL4AD-CIB1-tscADH1 | LEU2 | Ampicillin | AddGene #28245; Kennedy, et al 2010 |
| **pMM0160** | pGal4BD-CRY2 | pscADH1-GAL4BD-CRY2-tscADH1 | TRP1 | Kanamycin | AddGene #28243; Kennedy, et al 2010 |
| **pMM0162** | bSR97 | His3 Pfus1-yEVenus pSTL1-dsRED | HIS3 | Ampicillin | Ramanathan Lab, Unpublished |
| **pMM0223** | pKT90 | pFA6a–link–yEVenus–SpHIS5 | SpHIS5 | Ampicillin | Sheff, MA and KT Thorn 2004 Yeast |
| **pMM0281** | SV40NLS-VP16-CIB1 | pscADH1-SV40NLS-VP16-CIB1-tscADH1 | LEU2 | Ampcillin | Melendez, et al 2014 |
| **pMM0282** | pFLAG(3X)-NLS-ZIF268DBD-CRY2 (L10) TRP1 | pscADH1-FLAG(3X)-NLS-ZIF268DBD-CRY2 (L10)-tscADH1 TRP1 | TRP1 | Ampcillin | This study |
| **pMM0283** | pFLAG(3X)-NLS-ZIF268DBD-CRY2 (L4) TRP1 | pscADH1-FLAG(3X)-NLS-ZIF268DBD-CRY2 (L4)-tscADH1 TRP1 | TRP1 | Ampcillin | This study |
| **pMM0284** | FLAG(3X)-NLS-ZIF268DBD-CRY2 (L3) | pscADH1-pFLAG(3X)-NLS-ZIF268DBD-CRY2 (L3)-tscADH1 TRP1 | TRP1 | Ampcillin | This study |
| **pMM0285** | pZF(1BS) | pZF(1BS)-yEVENUS | URA3 | Ampcillin | This study |
| **pMM0286** | pZF(3oBS) | pZF(3oBS)-yEVENUS | URA3 | Ampcillin | This study |
| **pMM0287** | pZF(3BS) | pZF(3BS)-yEVENUS | URA3 | Ampcillin | This study |
| **pMM0288** | pZF(3BSc) | pZF(3BSc)-yEVENUS | URA3 | Ampcillin | This study |
| **pMM0289** | pZF(4BS) | pZF(4BSc)-yEVENUS | URA3 | Ampcillin | This study |
| **pMM0290** | pZF(3BSopp) | pZF(3BSop)-yEVENUS | URA3 | Ampcillin | This study |
| **pMM0296** | pSH65 | pGAL1-CRE PheloR | PHLEO | Ampicillin | Botstein Lab; Gueldener, et al 2002 |
| **pMM0301** | pGAL1-VENUS | pGAL1-yEVENUS CEN scURA3 | URA3 | Ampicillin | This study |
| **pMM0305** | pADH1(native)-FLAG(3X)-Zif268DBD-CRY2 | pADH1(native)-FLAG(3X)-Zif268DBD-CRY2 TRP1 CEN | TRP1 | Ampicillin | This study |
| **pMM0306** | pADH1_PacI-FLAG(3X)-Zif268DBD-CRY2 | pADH1_PacI-FLAG(3X)-Zif268DBD-CRY2 TRP1 CEN | TRP1 | Ampicillin | This study |
| **pMM0313** | SV40NLS-Zif268DBD-CRY2 (L3) | pscADH1-SV40NLS-Zif268DBD-CRY2 (L3)-tADH1 TRP CEN | TRP1 | Ampicillin | This study |
| **pMM0314** | H2BNLS-Zif268DBD-CRY2 (L3) | pscADH1-H2BNLS-Zif268DBD-CRY2 (L3)-tADH1 TRP CEN | TRP1 | Ampicillin | This study |
| **pMM0315** | pGal4AD | GAL4AD | LEU2 | Ampicillin | This study |
| **pMM0316** | pGal4BD | GAL4BD | TRP1 | Kanamycin | This study |
| **pMM0317** | NLS-ZIF268DBD-CRY2 (L3) | pscADH-SV40NLS-Zif268DBD-CRY2 (L3)-tscADH1 scTRP1 | TRP1 | Ampicillin | This study |
| **pMM0318** | FLAG(3X)-SV40NLS-Zif-ZCRY2-SV40NLS | pscADH-FLAG(3X)-SV40NLS-Zif268DBD-CRY2-SV40NLS-tscADH1 scTRP1 | TRP1 | Ampicillin | This study |
| **pMM0319** | FLAG(3X)-SV40NLS-Zif-ZCRYPHR | pscADH-FLAG(3X)-SV40NLS-Zif-ZCRYPHR(L3)-tscADH1 scTRP1 | TRP1 | Ampicillin | This study |

| | | | | | |
|---|---|---|---|---|---|
| **pMM0320** | SV40NLS-Zif268DBD-CRY2PHR | pscADH1-SV40NLS-ZIF268DBD-CRY2PHR (L3)-tscADH1 TRP1 | TRP1 | Ampicillin | This Study |
| **pMM0321** | FLAG(3X)-Zif268DBD-CRY2PHR-SV40NLS | pscADH-FLAG(3X)-SV40NLS-Zif-ZCRY2PHR-SV40NLS-tscADH1 scTRP1 | TRP1 | Ampicillin | This Study |
| **pMM0326** | pUG72 | loxP-KIURA3-loxP | NA | Ampicillin | Gueldener, *et al* 2002 [18] |
| **pMM0452** | pYTK001 | None | NA | Chloramphenicol | AddGene #65108; Lee, et al 2015 |
| **pMM0453** | pYTK009 | pscTDH3 | NA | Chloramphenicol | AddGene #65116; Lee, et al 2015 |
| **pMM0454** | pYTK017 | pscRPL18B | NA | Chloramphenicol | AddGene #65214; Lee, et al 2015 |
| **pMM0455** | pYTK027 | pscREV1 | NA | Chloramphenicol | AddGene #65134; Lee, et al 2015 |
| **pMM0456** | pYTK034 | mRUBY2 | NA | Chloramphenicol | AddGene #65141; Lee, et al 2015 |
| **pMM0457** | pYTK053 | tADH1 | NA | Chloramphenicol | AddGene #65160; Lee, et al 2015 |
| **pMM0458** | pYTK096 | 5' URA3 homology-ConLS'-GFP dropout-ConRE' URA3 URA3 3'-homology KanR-ColE1 | scURA3 | Kanamycin | Addgene #65203; Lee, et al 2015 |
| **pMM0477** | pYTK008 | ConLS' | NA | Chloramphenicol | Addgene #65115; Lee et al 2015 |
| **pMM0478** | pYTK073 | ConRE' | NA | Chloramphenicol | Addgene #65180; Lee, et al 2015 |
| **pMM0479** | pYTK075 | LEU2 | LEU2 | Chloramphenicol | AddGene #65182; Lee, et al 2015 |
| **pMM0480** | pYTK087 | LEU2 3' homology | NA | Chloramphenicol | AddGene #65194; Lee, et al 2015 |
| **pMM0481** | pYTK090 | KanR-ColE1 mRFP1 | NA | Kanamycin | AddGene #65197; Lee, et al 2015 |
| **pMM0482** | pYTK093 | LEU2 5' homology | NA | Chloramphenicol | AddGene #65200; Lee et al 2015 |
| **pMM0489** | pYTK002 | ConLS | NA | Chloramphenicol | AddGene #65109; Lee, et al 2015 |

| | | | | | |
|---|---|---|---|---|---|
| **pMM0490** | pYTK047 | sfGFP dropout | NA | Chloramphenicol | AddGene #65154; Lee, et al 2015 |
| **pMM0491** | pYTK067 | ConR1 | NA | Chloramphenicol | AddGene #65174; Lee, et al 2015 |
| **pMM0494** | yOTK_LEU2entry | LEU2 5' homology-ConLS-sfGFP dropout-ConR1-LEU2 3' homology | LEU2 | Kanamycin | This study |
| **pMM0495** | pTDH3-mRUBY2 | LEU2 5' homology-ConLS-pTDH3-mRUBY2-tADH1-ConR1-LEU2 3' homology | LEU2 | Kanamycin | This study |
| **pMM0496** | pRPL18B-mRUBY2 | LEU2 5' homology-ConLS-pRPL18B-mRUBY2-tADH1-ConR1-LEU2 3' homology | LEU2 | Kanamycin | This study |
| **pMM0497** | pREV1-mRUBY2 | LEU2 5' homology-ConLS-pREV1-mRUBY2-tADH1-ConR1-LEU2 3' homology | LEU2 | Kanamycin | This study |
| **pMM0515** | NLS-ZIF268DBD-CRY2 (L3) | SV40NLS-Zif268DBD-CRY2 (L3) CEN scTRP1 | TRP1 | Ampicillin | This study |
| **pMM0518** | pZF-3ZFBS-Venus | KanMXrev-pZF-3ZFBS-Venus | KanMX | Ampicillin | This study |
| **pMM0519** | loxP-KIURA3-loxP | loxP-KIURA3-loxP | KIURA3 | Chloramphenicol | This study |
| **pMM0520** | CIB1VP16 | SV40NLS-VP16-CIB1 LEU2 2µ | LEU2 | Ampicillin | This study |
| **pMM0521** | NLS-ZIF268DBD-CRY2 (L3) | SV40NLS-Zif268DBD-CRY2 (L3) scTRP1 | TRP1 | Ampicillin | This study |
| **pMM0522** | pYTK013 | pTEF1 | NA | Chloramphenicol | Addgene #65120; Lee, et al 2015 |
| **pMM0526** | pYTK086 | *URA3* 3' Homology | NA | Chloramphenicol | AddGene #65193; Lee, et al 2015 |
| **pMM0527** | pYTK092 | *URA3* 5' Homology | NA | Chloramphenicol | AddGene #65199; Lee, et al 2015 |
| **pMM0528** | pZF(3BS) | pZF(3BS) | NA | Chloramphenicol | This study |
| **pMM0529** | SV40NLS-ZiF268-CRY2 | SV40NLS-ZiF268DBD-CRY2 | NA | Chloramphenicol | This study |
| **pMM0530** | SV40NLS-ZiF268-CRY2PHR | SV40NLS-ZiF268DBD-CRY2PHR | NA | Chloramphenicol | This study |
| **pMM0531** | SV40NLS-VP16-CIB1 | SV40NLS-VP16-CIB1 | NA | Chloramphenicol | This study |
| **pMM0532** | pYTK003 | ConL1 | NA | Chloramphenicol | AddGene #65110; Lee, et al 2015 |
| **pMM0533** | pYTK004 | ConL2 | NA | Chloramphenicol | Addgene #65111; Lee, et al 2015 |
| **pMM0537** | pYTK068 | ConR2 | NA | Chloramphenicol | Addgene #65175; Lee, et al 2015 |
| **pMM0541** | pYTK072 | ConRE | NA | Chloramphenicol | Addgene #65179; Lee, et al 2015 |

| | | | | | |
|---|---|---|---|---|---|
| **pMM0542** | pYTK051 | tENO1 | NA | Chloramphenicol | Addgene #65158; Lee, et al 2015 |
| **pMM0543** | pYTK052 | tSSA1 | NA | Chloramphenicol | Addgene #65159; Lee, et al 2015 |
| **pMM0547** | pYTK048 | Spacer | NA | Chloramphenicol | Addgene Plasmid #65155; Lee, et al 2015 |
| **pMM0553** | pZF-mRUBY2 @ LEU2 | Leu2 5' homology-pZiF promoter-mRuby2-tADH1-Con1-LEU2-Leu2 3' homology | LEU2 | Kanamycin | This study |
| **pMM0554** | pTDH3-AD-CIB1 @ LEU2 | Leu2 5' homology-pTDH3-SV40NLS-VP16-CIB1-tADH1-Con1-LEU2-Leu2 3' homology | LEU2 | Kanamycin | This study |
| **pMM0556** | pYTK095 | sfGFP AmpR-ColE1 | NA | Ampicillin | AddGene #65202; Lee, et al 2015 |
| **pMM0557** | pTDH3-ZCRY2 @ LEU2 | Ura3 3' homology-pTDH3-SV40NLS-ZiF268-CRY2-tADH1-URA3-Ura3 5' homology | URA3 | Kanamycin | This study |
| **pMM0558** | pTHD3-ZCRY2PHR2 @ LEU2 | Ura3 3' homology-pTDH3-SV40NLS-ZiF268-CRY2PHR-tADH1-URA3-Ura3 5' homology | URA3 | Kanamycin | This study |
| **pMM0562** | pYTK023 | pscRNR2 | NA | Chloramphenicol | Addgene #65130; Lee, et al 2015 |
| **pMM0617** | pCS0010, A, loxP entry vector | KanR-ColE1 URA 5' homology-ConLS'-sfGFP dropout-ConRE'-loxP-KlURA3-loxP-URA3 5' homology | klURA3 | Kanamycin | This study |
| **pMM0619** | pCS0012, C, C3F spacer, background fluorescence control | ConL2-Spacer-ConRE-AmpR-Col-E1 | NA | Ampicillin | This study |
| **pMM0620** | pCS0013, D, C1_pTEF1_CIB1 | ConLS_pTEF1-SV40NLS-VP16-CIB1-tENO1-ConR1 AmpR Col-E1 | NA | Ampicillin | This study |
| **pMM0621** | pCS0014, E, C1_pRPL18B_CIB1 | ConLS_pRPL18B-SV40NLS-VP16-CIB1-tENO1-ConR1 AmpR Col-E1 | NA | Ampicillin | This study |
| **pMM0622** | pCS0015, F, C2_pTEF1_CRY2PHR | ConL1-pTEF1-SV40NLS-ZiF268DBD-CRY2PHR-tSSA1-ConR2 AmpR-ColE1 | NA | Ampicillin | This study |
| **pMM0623** | pCS0016, G, C2_pRPL18B_CRY2phr | ConL1-pRPL18B-SV40NLS-ZiF268DBD-CRY2PHR-tSSA1-ConR2 AmpR-ColE1 | NA | Ampicillin | This study |
| **pMM0624** | pCS0017, H, C3F_pZiF_mRuby2 | ConL2-pZF(BS)-mRUBY2-tADH1-ConRE AmpR-ColE1 | NA | Ampicillin | This study |
| **pMM0625** | pCS0018, I, C3F_pREV1_mRuby2 | ConL2-pREV1-mRUBY2-tADH1-ConRE AmpR-ColE1 | NA | Ampicillin | This study |
| **pMM0626** | pCS0019, J, C3F_pRPL18B_mRuby2 | ConL2-pRPL18B-mRUBY2-tADH1-ConRE AmpR-ColE1 | NA | Ampicillin | This study |
| **pMM0627** | pCS0020, K, C3F_pTDH3_mRuby2 | ConL2-pTDH3-mRUBY2-tADH1-ConRE AmpR-ColE1 | NA | Ampicillin | This study |
| **pMM0628** | pCS0021, L, C1_pRNR2_CIB1 | ConLS_pRNR2-SV40NLS-VP16-CIB1-tENO1-ConR1 AmpR Col-E1 | NA | Ampicillin | This study |
| **pMM0629** | pCS0022, M, C2_pRNR2_CRY2PHR | ConL1-pRNR2-SV40NLS-ZiF268DBD-CRY2PHR-tSSA1-ConR2 AmpR-ColE1 | NA | Ampicillin | This study |

| | | | | | |
|---|---|---|---|---|---|
| **pMM0637** | pCS0030, C1_pTEF1-CIB1 C2_pTEF1-CRY2PHR C3F_pZIF-mRuby2 | 5' URA3 homology-pTEF1-SV40NLS-VP16-CIB1-tENO1-pTEF1-SV40NLS-ZiF268DBD-CRY2PHR-tSSA1-pZF(BS)-mRUBY2-tADH1-URA3 URA3 3'-homology KanR-ColE1 | scURA 3 | Kanamycin | This study |
| **pMM0638** | pCS0031, C1_pTEF1-CIB1 C2_pRPL18B-CRY2PHR C3F_pZIF-mRuby2 | 5' URA3 homology-pTEF1-SV40NLS-VP16-CIB1-tENO1-pRPL18B-SV40NLS-ZiF268DBD-CRY2PHR-tSSA1-pZF(BS)-mRUBY2-tADH1-URA3 URA3 3'-homology KanR-ColE1 | scURA 3 | Kanamycin | This study |
| **pMM0639** | pCS0032, C1_pTEF1-CIB1 C2_pRNR2-CRY2PHR2 C3F_pZIF-mRuby2 | 5' URA3 homology-pTEF1-SV40NLS-VP16-CIB1-tENO1-pRNR2-SV40NLS-ZiF268DBD-CRY2PHR-tSSA1-pZF(BS)-mRUBY2-tADH1-URA3 URA3 3'-homology KanR-ColE1 | scURA 3 | Kanamycin | This study |
| **pMM0640** | pCS0033, C1_pRPL18B-CIB1 C2_pTEF1-CRY2PHR C3F_pZIF-mRuby2 | 5' URA3 homology-pRPL18B-SV40NLS-VP16-CIB1-tENO1-pTEF1-SV40NLS-ZiF268DBD-CRY2PHR-tSSA1-pZF(BS)-mRUBY2-tADH1-URA3 URA3 3'-homology KanR-ColE1 | scURA 3 | Kanamycin | This study |
| **pMM0641** | pCS0034, C1_pRPL18B-CIB1 C2_pRPL18B-CRY2PHR C3F_pZIF-mRuby2 | 5' URA3 homology-pRPL18B-SV40NLS-VP16-CIB1-tENO1-pRPL18B-SV40NLS-ZiF268DBD-CRY2PHR-tSSA1-pZF(BS)-mRUBY2-tADH1-URA3 URA3 3'-homology KanR-ColE1 | scURA 3 | Kanamycin | This study |
| **pMM0642** | pCS0035, C1_pRPL18B-CIB1 C2_pRNR2-CRY2PHR2 C3F_pZIF-mRuby2 | 5' URA3 homology-pRPL18B-SV40NLS-VP16-CIB1-tENO1-pRNR2-SV40NLS-ZiF268DBD-CRY2PHR-tSSA1-pZF(BS)-mRUBY2-tADH1-URA3 URA3 3'-homology KanR-ColE1 | scURA 3 | Kanamycin | This study |
| **pMM0643** | pCS0036, C1_pRNR2-CIB1 C2_pTEF1-CRY2PHR C3F_pZIF-mRuby2 | 5' URA3 homology-pRNR2-SV40NLS-VP16-CIB1-tENO1-pTEF1-SV40NLS-ZiF268DBD-CRY2PHR-tSSA1-pZF(BS)-mRUBY2-tADH1-URA3 URA3 3'-homology KanR-ColE1 | scURA 3 | Kanamycin | This study |
| **pMM0644** | pCS0037, C1_pRNR2-CIB1 C2_pRPL18B-CRY2PHR C3F_pZIF-mRuby2 | 5' URA3 homology-pRNR2-SV40NLS-VP16-CIB1-tENO1-pRPL18B-SV40NLS-ZiF268DBD-CRY2PHR-tSSA1-pZF(BS)-mRUBY2-tADH1-URA3 URA3 3'-homology KanR-ColE1 | scURA 3 | Kanamycin | This study |
| **pMM0645** | pCS0038, C1_pRNR2-CIB1 C2_pRNR2-CRY2-PHR2 C3F_pZIF-mRuby2 | 5' URA3 homology-pRNR2-SV40NLS-VP16-CIB1-tENO1-pRNR2-SV40NLS-ZiF268DBD-CRY2PHR-tSSA1-pZF(BS)-mRUBY2-tADH1-URA3 URA3 3'-homology KanR-ColE1 | scURA 3 | Kanamycin | This study |
| **pMM0646** | pCS0040, C1_pTEF1-CIB1 C2_pRPL18B-CRY2PHR C3F_pZIF-mRuby2 | KanR-ColE1 URA 5' homology-pTEF1-SV40NLS-VP16-CIB1-tENO1--pRPL18B-SV40NLS-ZiF268DBD-CRY2PHR-tSSA1--pZF(BS)-mRUBY2-tADH1--loxP-KIURA3-loxP-URA3 3' homology | KIURA3 | Kanamycin | This study |
| **pMM0647** | pCS0043, C1_pRPL18B-CIB1 C2_pRPL18B-CRY2PHR C3F_pZIF-mRuby2 | KanR-ColE1 URA 5' homology-pRPL18B-SV40NLS-VP16-CIB1-tENO1--pRPL18B-SV40NLS-ZiF268DBD-CRY2PHR-tSSA1--pZF(BS)-mRUBY2-tADH1--loxP-KIURA3-loxP-URA3 3' homology | KIURA3 | Kanamycin | This study |
| **pMM0648** | pCS0048, C1_pTEF1-CIB1 C2_pRPL18B-CRY2PHR C3F_spacer | 5' URA3 homology-pTEF1-SV40NLS-VP16-CIB1-tENO1-pRPL18B-SV40NLS-ZiF268DBD-CRY2PHR-tSSA1-Spacer-URA3 URA3 3'-homology KanR-ColE1 | scURA 3 | Kanamycin | This study |
| **pMM0649** | pCS0049, C1_pTEF1-CIB1 C2_pRPL18B-CRY2PHR C3F_pREV1-mRuby2 | 5' URA3 homology-pTEF1-SV40NLS-VP16-CIB1-tENO1-pRPL18B-SV40NLS-ZiF268DBD-CRY2PHR-tSSA1-pREV1-mRUBY2-tADH1-URA3 URA3 3'-homology KanR-ColE1 | scURA 3 | Kanamycin | This study |
| **pMM0650** | pCS0050, C1_pTEF1-CIB1 C2_pRPL18B-CRY2PHR C3F_pRPL18B-mRuby2 | 5' URA3 homology-pTEF1-SV40NLS-VP16-CIB1-tENO1-pRPL18B-SV40NLS-ZiF268DBD-CRY2PHR-tSSA1-pRPL18B-mRUBY2-tADH1-URA3 URA3 3'-homology KanR-ColE1 | scURA 3 | Kanamycin | This study |
| **pMM0651** | pCS0051, C1_pTEF1-CIB1 C2_pRPL18B-CRY2PHR C3F_pTDH3-mRuby2 | 5' URA3 homology-pTEF1-SV40NLS-VP16-CIB1-tENO1-pRPL18B-SV40NLS-ZiF268DBD-CRY2PHR-tSSA1-pTDH3-mRUBY2-tADH1-URA3 URA3 3'-homology KanR-ColE1 | scURA 3 | Kanamycin | This study |
| **pMM0652** | pCS0052, C1_pRPL18B-CIB1 | 5' URA3 homology-pRPL18B-SV40NLS-VP16-CIB1-tENO1-pRPL18B-SV40NLS-ZiF268DBD-CRY2PHR-tSSA1-Spacer-URA3 URA3 3'-homology KanR-ColE1 | scURA 3 | Kanamycin | This study |

| | | | | | |
|---|---|---|---|---|---|
| | C2_pRPL18B-CRY2PHR C3F_spacer | | | | |
| **pMM0653** | pCS0053, C1_pRPL18B-CIB1 C2_pRPL18B-CRY2PHR C3F_pREV1-mRuby2 | 5' URA3 homology-pRPL18B-SV40NLS-VP16-CIB1-tENO1-pRPL18B-SV40NLS-ZiF268DBD-CRY2PHR-tSSA1-pREV1-mRUBY2-tADH1-URA3 URA3 3'-homology KanR-ColE1 | scURA 3 | Kanamycin | This study |
| **pMM0654** | pCS0054, C1_pRPL18B-CIB1 C2_pRPL18B-CRY2PHR C3F_pRPL19B-mRuby2 | 5' URA3 homology-pRPL18B-SV40NLS-VP16-CIB1-tENO1-pRPL18B-SV40NLS-ZiF268DBD-CRY2PHR-tSSA1-pRPL18B-mRUBY2-tADH1-URA3 URA3 3'-homology KanR-ColE1 | scURA 3 | Kanamycin | This study |
| **pMM0655** | pCS0055, C1_pRPL18B-CIB1 C2_pRPL18B-CRY2PHR C3F_pTDH3-mRuby2 | 5' URA3 homology-pRPL18B-SV40NLS-VP16-CIB1-tENO1-pRPL18B-SV40NLS-ZiF268DBD-CRY2PHR-tSSA1-pTDH3-mRUBY2-tADH1-URA3 URA3 3'-homology KanR-ColE1 | scURA 3 | Kanamycin | This study |
| **pMM0656** | pCS0056 | ConL1-pTEF1-SV40NLS-ZiF268DBD-CRY2-tSSA1-ConR2-ColE1-AmpR | NA | Ampicillin | This study |
| **pMM0657** | pCS0057 | ConL1-pRPL18B-SV40NLS-ZiF268DBD-CRY2-tSSA1-ConR2-ColE1-AmpR | NA | Ampicillin | This study |
| **pMM0658** | pCS0058 | ConL1-pRNR2-SV40NLS-ZiF268DBD-CRY2-tSSA1-ConR2-ColE1-AmpR | NA | Ampicillin | This study |

Supplemental Figure 1

```
                10        20        30        40        50        60        70        80        90
       ....|....|....|....|....|....|....|....|....|....|....|....|....|....|....|....|....|....|....
PGAL1  TTATATTGAATTTTCAAAAATTCTTACTTTTTTTTTGGATGGACGCAAAGAAGTTTAATAATCATATTACATGGCATTACCACCATATACATAT
1BS    TTATATTGAATTTTCAAAAATTCTTACTTTTTTTTTGGATGGACGCAAAGAAGTTTAATAATCATATTACATGGCATTACCACCATATACATAT
2BS    TTATATTGAATTTTCAAAAATTCTTACTTTTTTTTTGGATGGACGCAAAGAAGTTTAATAATCATATTACATGGCATTACCACCATATACATAT
3BS    TTATATTGAATTTTCAAAAATTCTTACTTTTTTTTTGGATGGACGCAAAGAAGTTTAATAATCATATTACATGGCATTACCACCATATACATAT
3BS*   TTATATTGAATTTTCAAAAATTCTTACTTTTTTTTTGGATGGACGCAAAGAAGTTTAATAATCATATTACATGGCATTACCACCATATACATAT
4BS*   TTATATTGAATTTTCAAAAATTCTTACTTTTTTTTTGGATGGACGCAAAGAAGTTTAATAATCATATTACATGGCATTACCACCATATACATAT
3BSopp TTATATTGAATTTTCAAAAATTCTTACTTTTTTTTTGGATGGACGCAAAGAAGTTTAATAATCATATTACATGGCATTACCACCATATACATAT

               110       120       130       140       150       160       170       180       190
       ....|....|....|....|....|....|....|....|....|....|....|....|....|....|....|....|....|....|....
PGAL1  ACATATCCATATCTAATCTTACTTATATGTTGTGGAAATGTAAAGAGCCCCATTATCTTAGCCTAAAAAAACCTTCTCTTTGGAACTTTCAGTA
1BS    ACATATCCATATCTAATCTTACTTATATGTTGTGGAAATGTAAAGAGCCCCATTATCTTAGCCTAAAAAAACCTTCTCTTTGGAACTTTCAGTA
2BS    ACATATCCATATCTAATCTTACTTATATGTTGTGGAAATGTAAAGAGCCCCATTATCTTAGCCTAAAAAAACCTTCTCTTTGGAACTTTCAGTA
3BS    ACATATCCATATCTAATCTTACTTATATGTTGTGGAAATGTAAAGAGCCCCATTATCTTAGCCTAAAAAAACCTTCTCTTTGGAACTTTCAGTA
3BS*   ACATATCCATATCTAATCTTACTTATATGTTGTGGAAATGTAAAGAGCCCCATTATCTTAGCCTAAAAAAACCTTCTCTTTGGAACTTTCAGTA
4BS*   ACATATCCATATCTAATCTTACTTATATGTTGTGGAAATGTAAAGAGCCCCATTATCTTAGCCTAAAAAAACCTTCTCTTTGGAACTTTCAGTA
3BSopp ACATATCCATATCTAATCTTACTTATATGTTGTGGAAATGTAAAGAGCCCCATTATCTTAGCCTAAAAAAACCTTCTCTTTGGAACTTTCAGTA

               210       220       230       240       250       260       270       280       290
       ....|....|....|....|....|....|....|....|....|....|....|....|....|....|....|....|....|....|....
PGAL1  TTAACTGCTCATTGCTATATTGAAGTGCGGCCGCGAAGCCGCCGAGCGGGCGACAGCCCTCCGACGG----AAGACTCTAGACCGTGCGTCCTC
1BS    TTAACTGCTCATTGCTATATTGAAGTGCGGCCGCG---CGTGGGCGT------------------------------CTAGACCGTGCGTCCTC
2BS    TTAACTGCTCATTGCTATATTGAAGTGCGGCCGCG---CGTGGGCGTGGGCGG-----CGTGGGCGT----------CTAGACCGTGCGTCCTC
3BS    TTAACTGCTCATTGCTATATTGAAGTGCGGCCGCG---CGTGGGCGAGCGTGGGCGAGCGTGGGCGT----------CTAGACCGTGCGTCCTC
3BS*   TTAACTGCTCATTGCTATATTGAAGTGCGGCCGCG---CGTGGGCGTGCGTGGGCGTGCGTGGGCGT----------CTAGACCGTGCGTCCTC
4BS*   TTAACTGCTCATTGCTATATTGAAGTGCGGCCGCG---CGTGGGCGTGCGTGGGCGTGCGTGGGCGTGCGTGGGCGTCTAGACCGTGCGTCCTC
3BSopp TTAACTGCTCATTGCTATATTGAAGTGCGGCCGC----CGCCCACGCACGCCCACGCACGCCCACG---------CTCTAGACCGTGCGTCCTC

               310       320       330       340       350       360       370       380       390
       ....|....|....|....|....|....|....|....|....|....|....|....|....|....|....|....|....|....|....
PGAL1  ACCGGTCGCGTTCCTGAAACGCAGATGTGCCTCGCGCCGCACTGCTCCGAACAATAAAGATTCTACAATACTAGCTTTTATGGTTATGAAGAGG
1BS    ACCGGTCGCGTTCCTGAAACGCAGATGTGCCTCGCGCCGCACTGCTCCGAACAATAAAGATTCTACAATACTAGCTTTTATGGTTATGAAGAGG
2BS    ACCGGTCGCGTTCCTGAAACGCAGATGTGCCTCGCGCCGCACTGCTCCGAACAATAAAGATTCTACAATACTAGCTTTTATGGTTATGAAGAGG
3BS    ACCGGTCGCGTTCCTGAAACGCAGATGTGCCTCGCGCCGCACTGCTCCGAACAATAAAGATTCTACAATACTAGCTTTTATGGTTATGAAGAGG
3BS*   ACCGGTCGCGTTCCTGAAACGCAGATGTGCCTCGCGCCGCACTGCTCCGAACAATAAAGATTCTACAATACTAGCTTTTATGGTTATGAAGAGG
4BS*   ACCGGTCGCGTTCCTGAAACGCAGATGTGCCTCGCGCCGCACTGCTCCGAACAATAAAGATTCTACAATACTAGCTTTTATGGTTATGAAGAGG
3BSopp ACCGGTCGCGTTCCTGAAACGCAGATGTGCCTCGCGCCGCACTGCTCCGAACAATAAAGATTCTACAATACTAGCTTTTATGGTTATGAAGAGG

               410       420       430       440       450       460       470       480       490
       ....|....|....|....|....|....|....|....|....|....|....|....|....|....|....|....|....|....|....
PGAL1  TGGCAGTAACCTGGCCCCACAAACCTTCAAATTAACGAATCAAATTAACAACCATAGGATGATAATGCGATTAGTTTTTTAGCCTTATTTCTGG
1BS    TGGCAGTAACCTGGCCCCACAAACCTTCAAATTAACGAATCAAATTAACAACCATAGGATGATAATGCGATTAGTTTTTTAGCCTTATTTCTGG
2BS    TGGCAGTAACCTGGCCCCACAAACCTTCAAATTAACGAATCAAATTAACAACCATAGGATGATAATGCGATTAGTTTTTTAGCCTTATTTCTGG
3BS    TGGCAGTAACCTGGCCCCACAAACCTTCAAATTAACGAATCAAATTAACAACCATAGGATGATAATGCGATTAGTTTTTTAGCCTTATTTCTGG
3BS*   TGGCAGTAACCTGGCCCCACAAACCTTCAAATTAACGAATCAAATTAACAACCATAGGATGATAATGCGATTAGTTTTTTAGCCTTATTTCTGG
4BS*   TGGCAGTAACCTGGCCCCACAAACCTTCAAATTAACGAATCAAATTAACAACCATAGGATGATAATGCGATTAGTTTTTTAGCCTTATTTCTGG
3BSopp TGGCAGTAACCTGGCCCCACAAACCTTCAAATTAACGAATCAAATTAACAACCATAGGATGATAATGCGATTAGTTTTTTAGCCTTATTTCTGG

               510       520       530       540       550       560       570       580       590
       ....|....|....|....|....|....|....|....|....|....|....|....|....|....|....|....|....|....|....
PGAL1  TAATCAGCGAAGCGATGATTTTTGATCTATTAACAGATATATAAATGGAAAAGCTGCATAACCACTTTAACTAATACTTTCAACATTTTCAGTT
1BS    TAATCAGCGAAGCGATGATTTTTGATCTATTAACAGATATATAAATGGAAAAGCTGCATAACCACTTTAACTAATACTTTCAACATTTTCAGTT
2BS    TAATCAGCGAAGCGATGATTTTTGATCTATTAACAGATATATAAATGGAAAAGCTGCATAACCACTTTAACTAATACTTTCAACATTTTCAGTT
3BS    TAATCAGCGAAGCGATGATTTTTGATCTATTAACAGATATATAAATGGAAAAGCTGCATAACCACTTTAACTAATACTTTCAACATTTTCAGTT
3BS*   TAATCAGCGAAGCGATGATTTTTGATCTATTAACAGATATATAAATGGAAAAGCTGCATAACCACTTTAACTAATACTTTCAACATTTTCAGTT
4BS*   TAATCAGCGAAGCGATGATTTTTGATCTATTAACAGATATATAAATGGAAAAGCTGCATAACCACTTTAACTAATACTTTCAACATTTTCAGTT
3BSopp TAATCAGCGAAGCGATGATTTTTGATCTATTAACAGATATATAAATGGAAAAGCTGCATAACCACTTTAACTAATACTTTCAACATTTTCAGTT

               610       620       630       640       650       660       670       680       690
       ....|....|....|....|....|....|....|....|....|....|....|....|....|....|....|....|....|....|....
PGAL1  ACTTCTTATTCAAATGTCATAAAAGTATCAACAAAAAATTGTTAATATACCTCTATACTTTAACGTCAAGGAGAAAAAACTATACTCGAGGTCG
1BS    ACTTCTTATTCAAATGTCATAAAAGTATCAACAAAAAATTGTTAATATACCTCTATACTTTAACGTCAAGGAGAAAAAACTATACTCGAGGTCG
2BS    ACTTCTTATTCAAATGTCATAAAAGTATCAACAAAAAATTGTTAATATACCTCTATACTTTAACGTCAAGGAGAAAAAACTATACTCGAGGTCG
3BS    ACTTCTTATTCAAATGTCATAAAAGTATCAACAAAAAATTGTTAATATACCTCTATACTTTAACGTCAAGGAGAAAAAACTATACTCGAGGTCG
3BS*   ACTTCTTATTCAAATGTCATAAAAGTATCAACAAAAAATTGTTAATATACCTCTATACTTTAACGTCAAGGAGAAAAAACTATACTCGAGGTCG
4BS*   ACTTCTTATTCAAATGTCATAAAAGTATCAACAAAAAATTGTTAATATACCTCTATACTTTAACGTCAAGGAGAAAAAACTATACTCGAGGTCG
3BSopp ACTTCTTATTCAAATGTCATAAAAGTATCAACAAAAAATTGTTAATATACCTCTATACTTTAACGTCAAGGAGAAAAAACTATACTCGAGGTCG

               710       720       730
       ....|....|....|....|....|....|
PGAL1  TCGATAAGCTTGATATCGAATTCCTGCAGCCCGGG
1BS    TCGATAAGCTTGATATCGAATTCCTGCAGCCCGGG
2BS    TCGATAAGCTTGATATCGAATTCCTGCAGCCCGGG
3BS    TCGATAAGCTTGATATCGAATTCCTGCAGCCCGGG
3BS*   TCGATAAGCTTGATATCGAATTCCTGCAGCCCGGG
4BS*   TCGATAAGCTTGATATCGAATTCCTGCAGCCCGGG
3BSopp TCGATAAGCTTGATATCGAATTCCTGCAGCCCGGG
```

Supplemental Figure 2

a

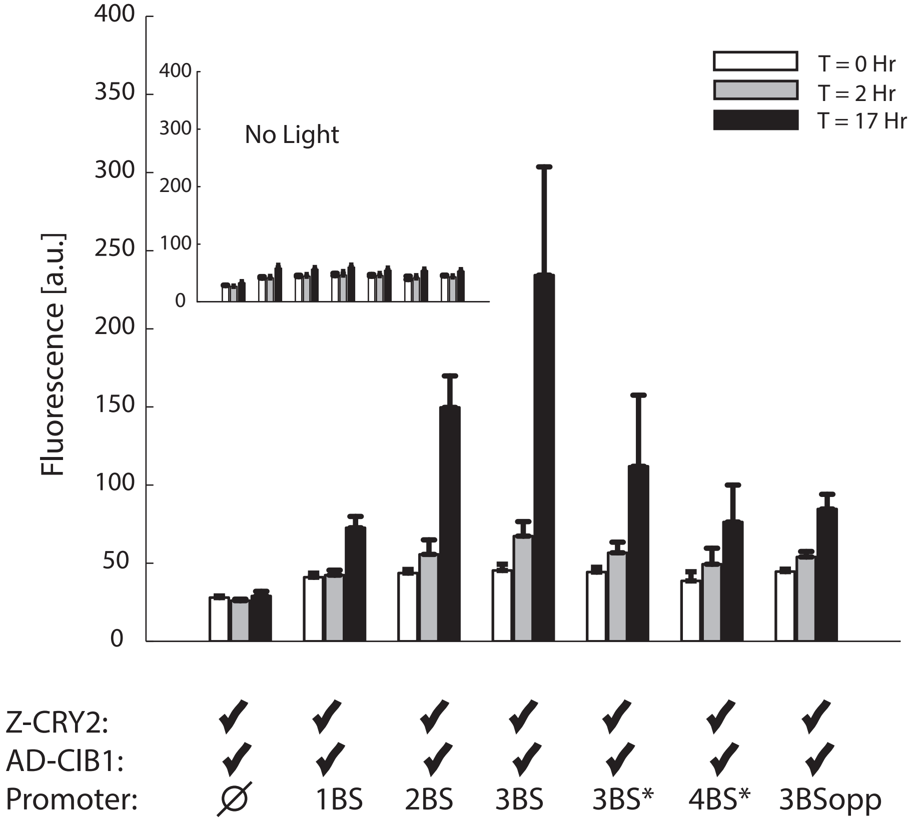

Lorem ipsum

b

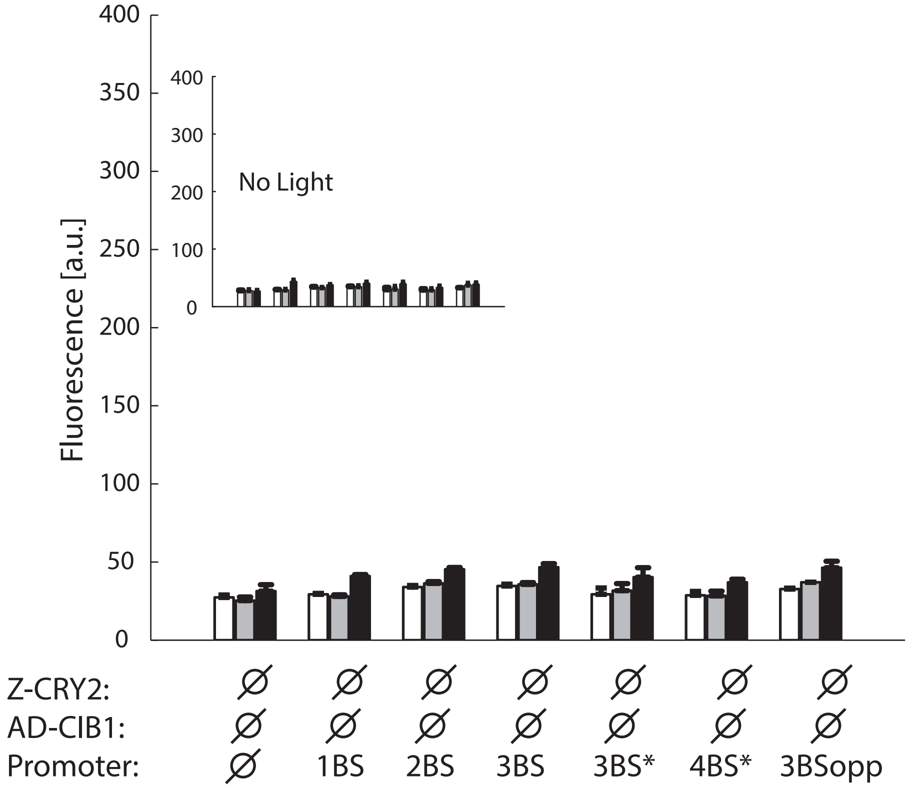

Supplemental Figure 3

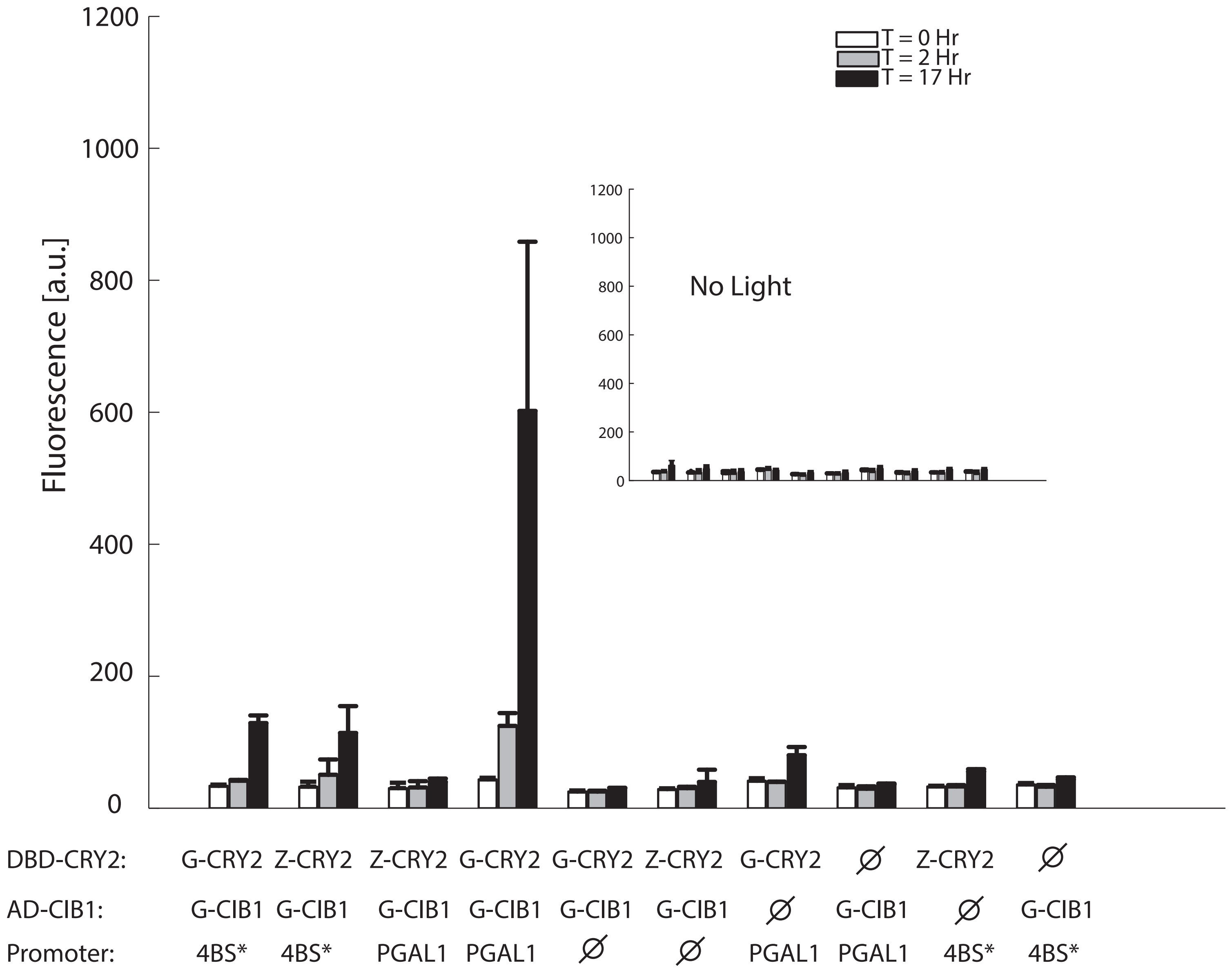

Supplemental Figure 4

a

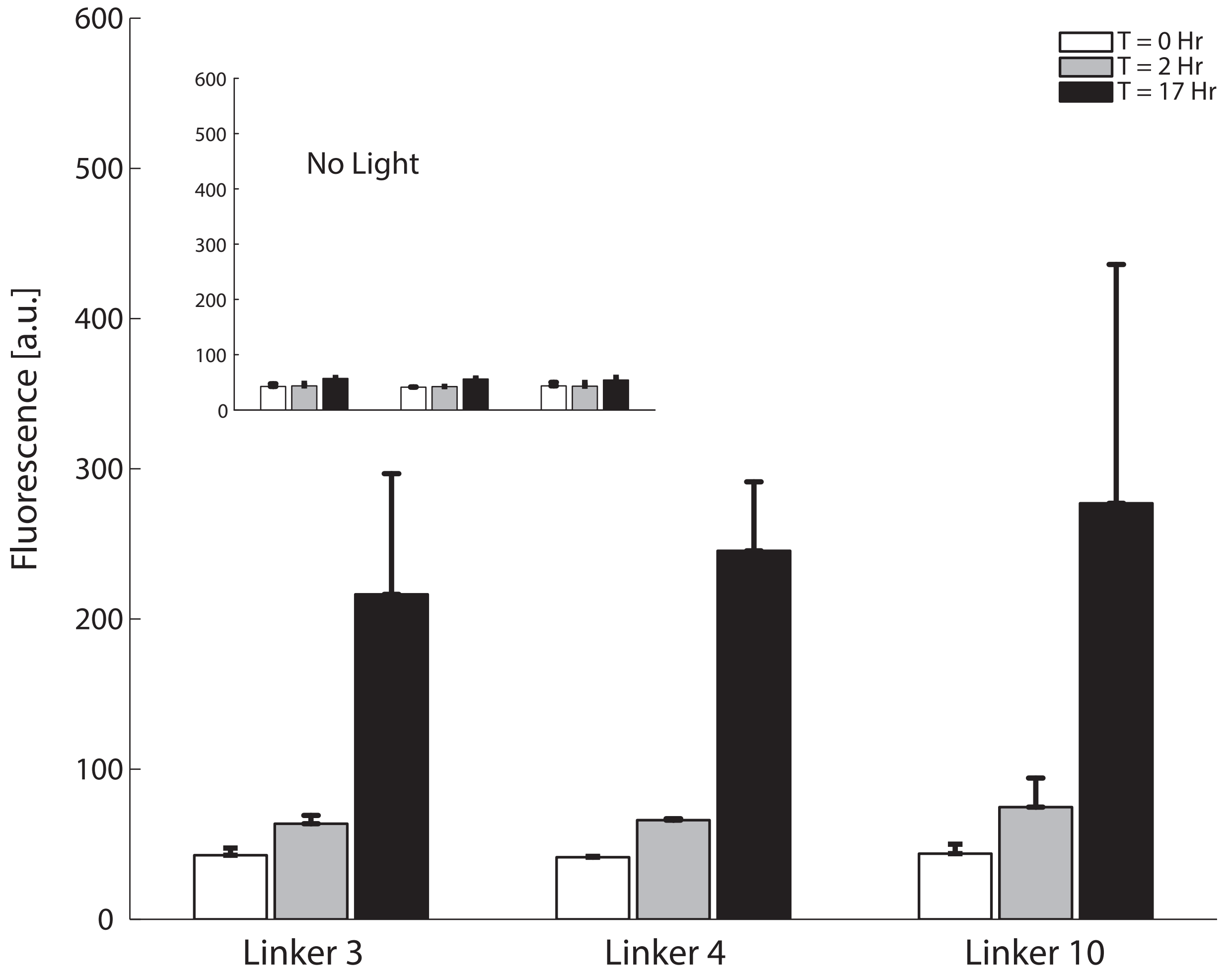

b

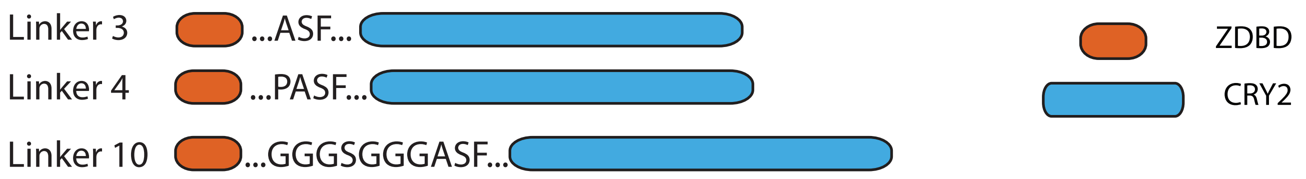

Supplemental Figure 5

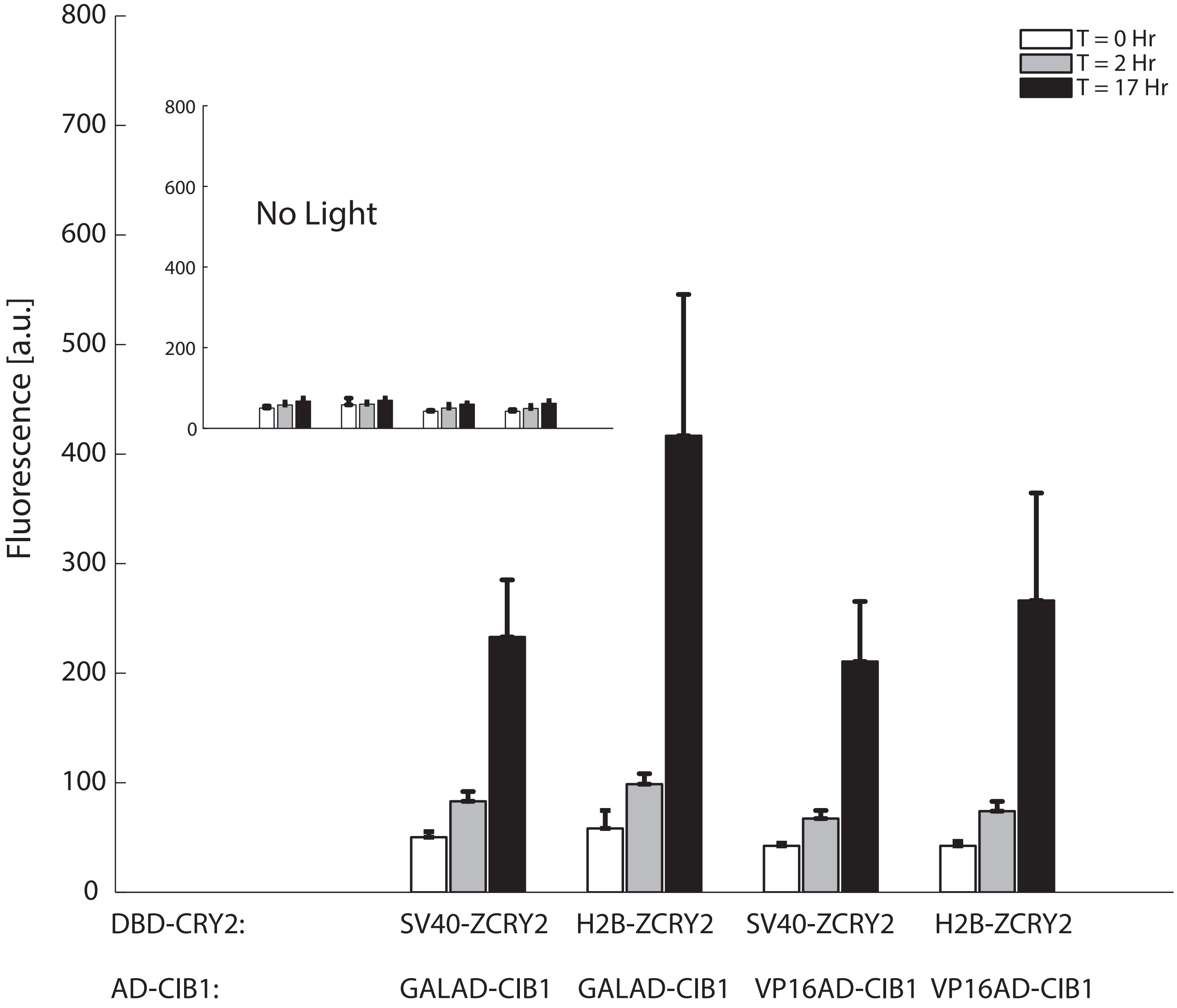

### Supplemental Figure 6

A

| Assembly Connector 1 | Promoter 2 | Coding Sequences 3 | Terminator 4 | Assembly Connector 5 | S. cerevisiae marker 6 |
|---|---|---|---|---|---|
| ConLS | pTEF1 | **AD-CIB1** | tENO1 | ConR1 | URA3 |
| ConL1 | pRPL18B | **ZDBD-CRY2PHR** | tSSA1 | ConR2 | **loxP-kiURA3-loxP** |
| ConL2 | pRNR2 | **ZDBD-CRY2** | tADH1 | ConRE | |
| ConLS' | **pZF(3BS)** | mRuby2 | | ConRE' | |
| | pREV1 | | | | |
| | pTDH3 | | | | |

B

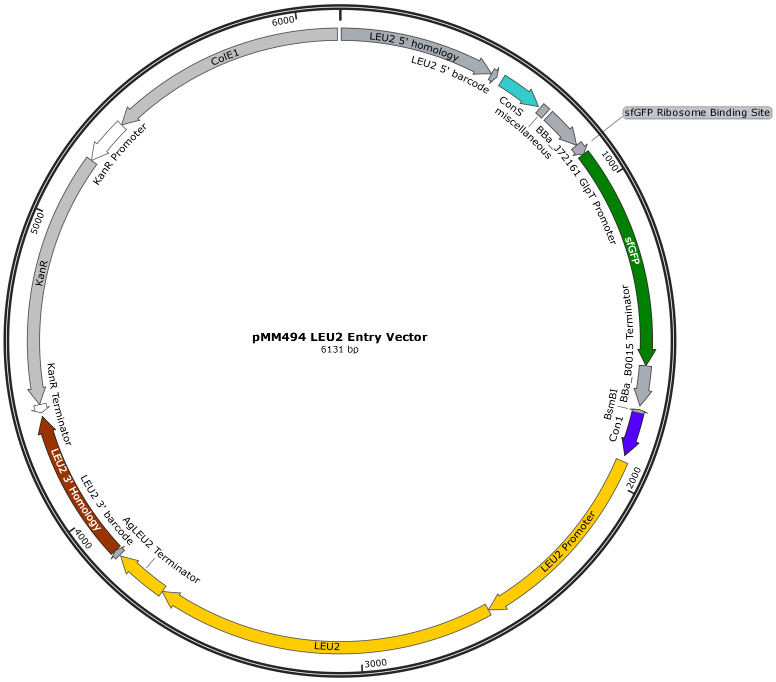

C

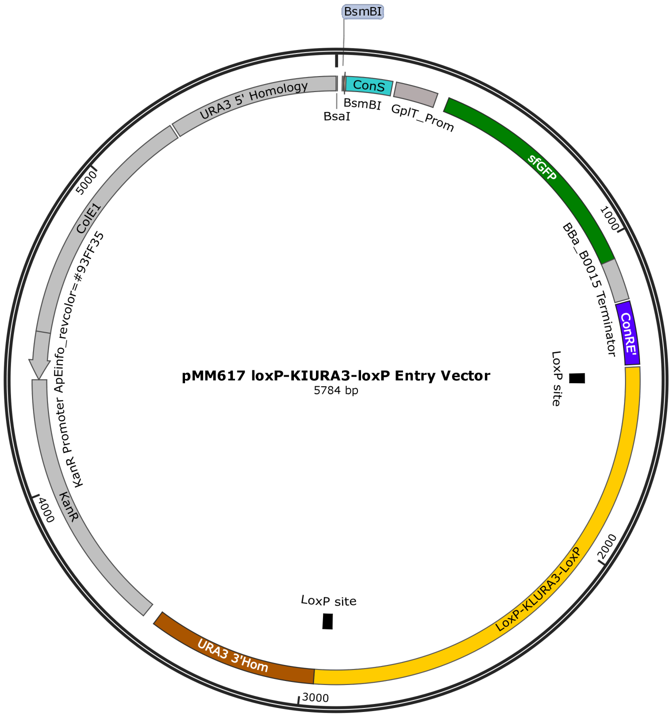

### Supplemental Figure 7

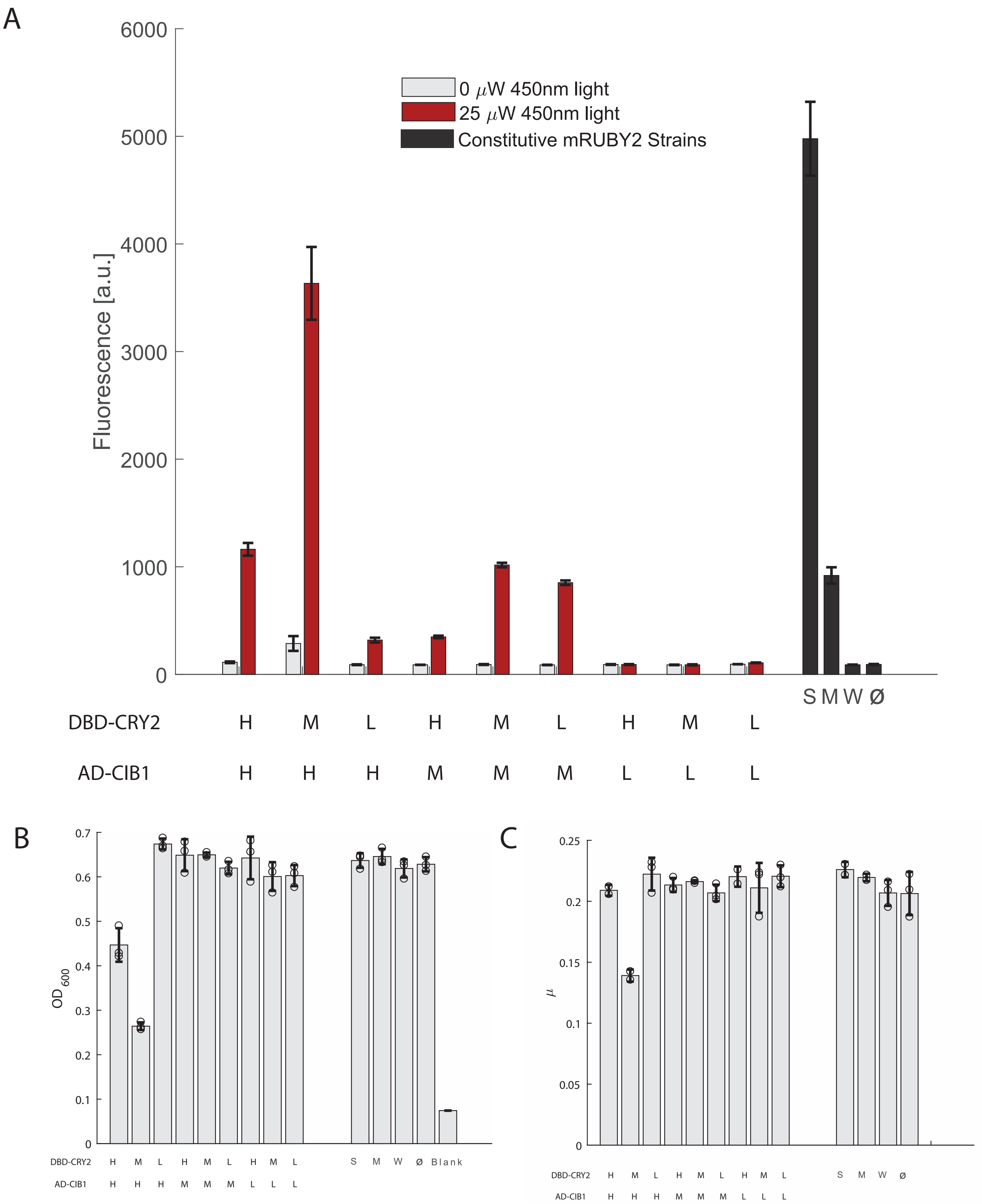

### Supplemental Figure 8

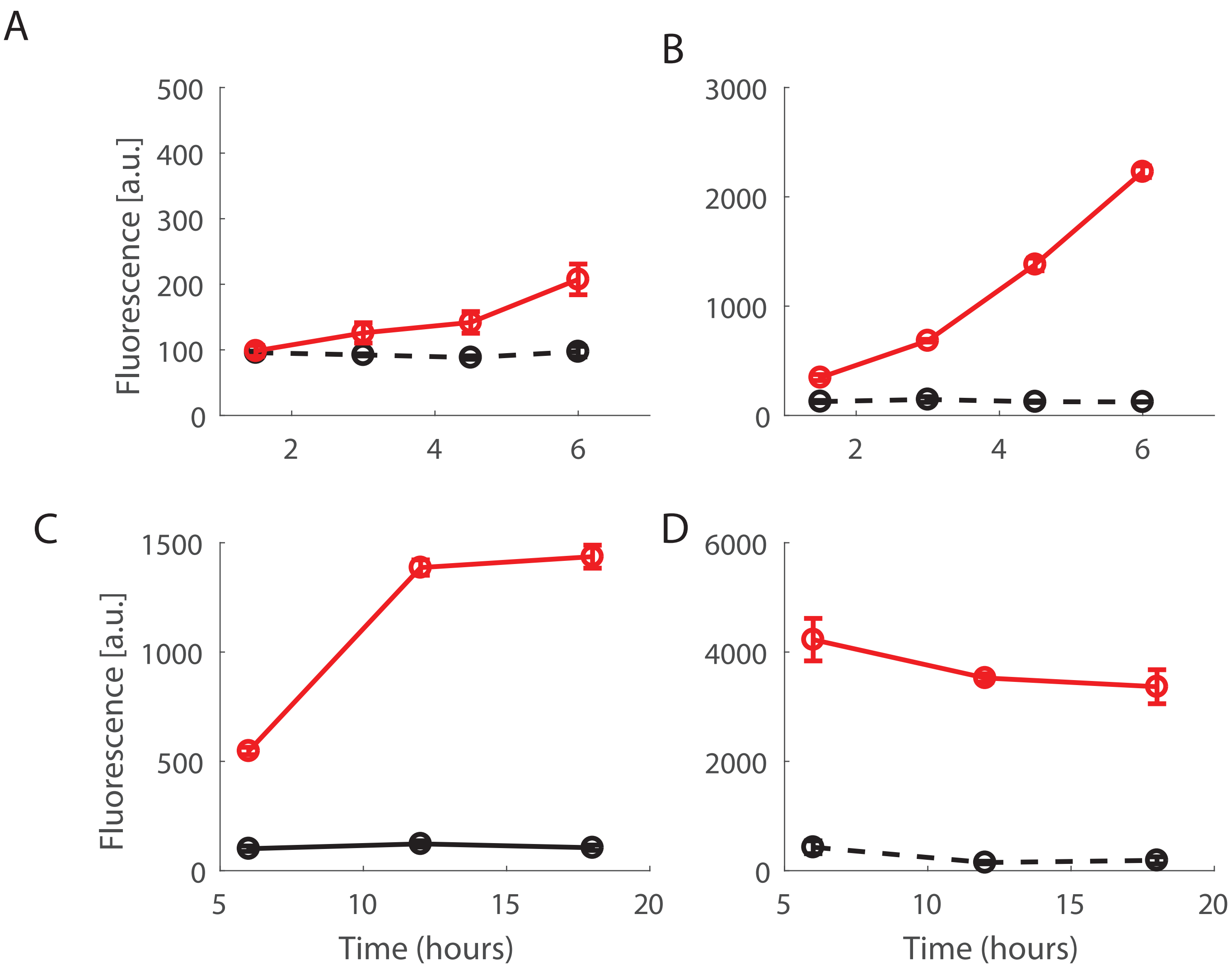

Supplemental Figure 9

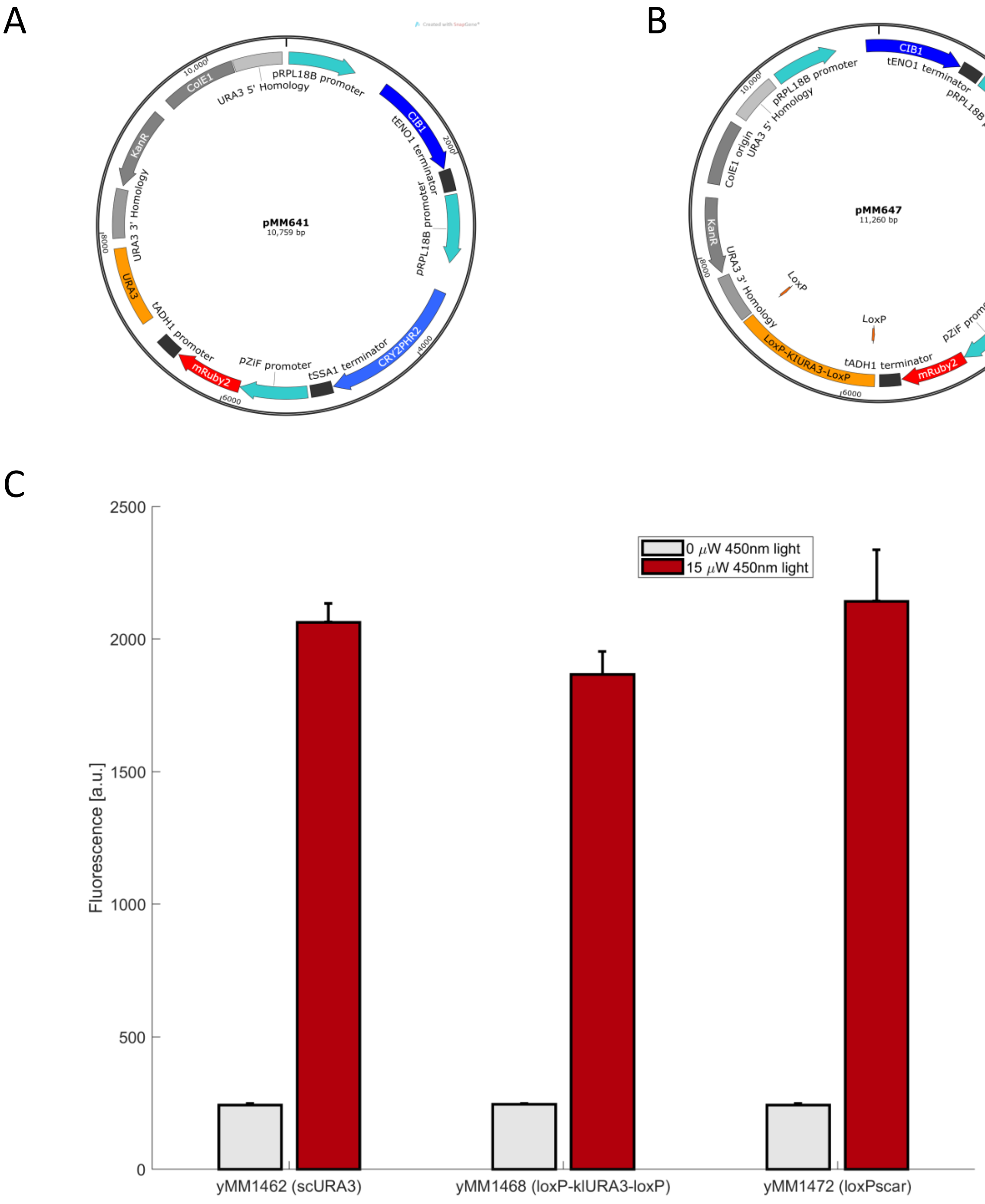
